# Anisotropic Hydrogel Degradation Enhances 3D Collective Mesenchymal Stromal Cell Alignment, Mechanotransduction and Osteogenic Differentiation

**DOI:** 10.1101/2024.04.11.588803

**Authors:** Claudia A. Garrido, Daniela S. Garske, Christian H. Bucher, Shahrouz Amini, Georg N. Duda, Katharina Schmidt-Bleek, Amaia Cipitria

## Abstract

Tissue engineering involves assembling cells and mimicking the complex anisotropic architecture of biological tissues to perform specific functions. This study explores the impact of patterned hydrogels with anisotropic degradation compared to single-phase materials on the potential of enhancing cell spreading, collective alignment, mechanotransduction and osteogenic differentiation of encapsulated human mesenchymal stromal cells (hMSCs). Spatial patterns of degradable (Deg) and non-degradable (noDeg) subregions are formed by photolithography: UV-triggered thiol-ene crosslinking with matrix metalloprotease (MMP) sensitive peptides form Deg phases, while non-UV exposed regions result in Diels-Alder spontaneous click crosslinking and noDeg phases. 3D patterns in hydrogel degradation enhance hMSC spreading and allow collective cell alignment in Deg areas, while cells remain rounded with no alignment in noDeg regions. In addition, we observe a boosted osteogenic differentiation when compared to single-phase materials, as mid osteogenic markers (osteocalcin) are expressed at day 14 in anisotropic gels, whereas in single-phase only early osteogenic markers are found (osterix). Mechanosensing pathways were evaluated using the expression and localization of YAP. Deg sections in patterned materials have an enhanced nuclear translocation and higher YAP expression compared to single-phase Deg materials and noDeg sections. This effect is lost and no patterns in YAP expression and localization emerge when using an MMP-scramble peptide or no-RGD materials. These findings demonstrate that 3D patterns in alginate hydrogel degradation guide hMSC spreading, collective alignment, enhance YAP nuclear translocation and osteogenic differentiation. Mimicking tissue anisotropy in 3D patterned hydrogels could have broad applications in biofabrication and tissue engineering.

Graphical Abstract

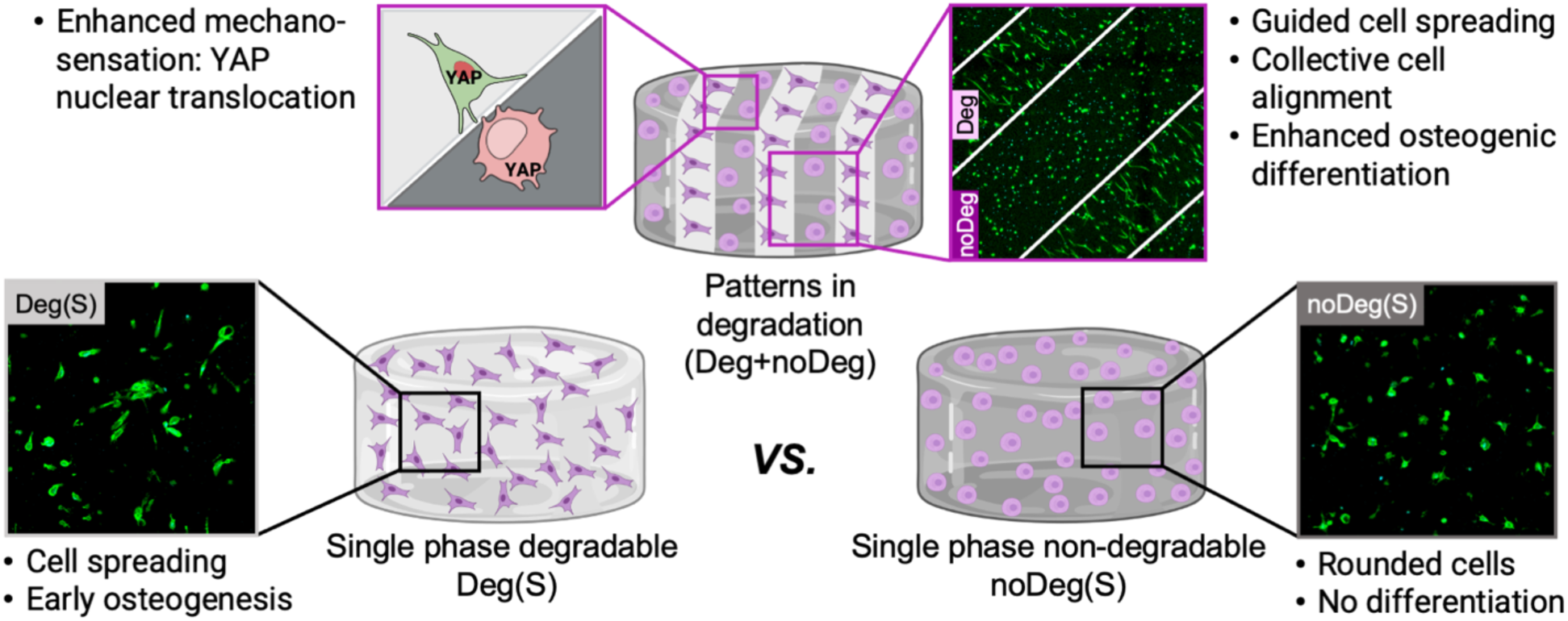

## 1 Introduction

Tissue engineering involves assembling cells and mimicking the complex anisotropic architecture of biological tissues to perform specific functions. The use of human mesenchymal stromal cells (hMSCs) is a promising approach in regenerative medicine [1], due to their self-renewal, clonality (proliferation from a single cell) and potency (differentiation into different cell types) [2].

Encapsulation of hMSCs in biomaterials offers a variety of options to guide and control their behavior, by modulating the local chemistry and physics provided to cells [3][4]. Hydrogels are widely used biomaterials thanks to their high cytocompatibility [5] and modifiable chemistry, providing tunable biophysical [6] and biochemical [7] properties. These have been shown to trigger specific cell activities, such as matrix stiffness [8][9][10], viscoelasticity [11] [12], degradability [13], ECM interaction by adhesion ligands [14], or ECM fiber arrangements [15]. In many cases these efforts have focused on homogeneous materials in which the cells reside. However, biomaterials in nature are typically anisotropic and provide a distinct architecture to cells.

Thus, single-phase materials can only partially provide a mimicry of the complex ECM interaction in living tissues. Native ECM is anisotropic with distinct spatially segregated properties [16]. Multiple phases in one matrix arranged in spatially discrete sections - patterned biomaterials - allow to mimic this natural ECM anisotropy, while keeping the benefits of a well-controlled synthetic material niche. The versatility of 3D patterned materials allows to tune biophysical properties and guide cell behavior. For example, patterns in biomolecule presentation have been used to spatially guide cell migration and alignment [17][18]. Despite their potential, the study of 3D patterned synthetic niches as analogs of anisotropic tissue architecture is still limited, especially how patterns in degradation can influence cell behavior. Degradable patterned microchannels have been used to guide endothelial cell sprouting [19], endothelium formation [20] and cell invasion [21] [22]. For other cell types, such as fibroblasts [23] or hMSCs [24], materials with patterns in degradation and stiffness have been studied in 3D. However, the effect of patterned matrix degradation, without the influence of other biophysical/biochemical factors, in controlling encapsulated cell behavior in multicompartment materials has not been fully elucidated yet.

The mechanisms underlying cell-matrix interactions in 3D degradable materials are traction forces [25] that interact with the dynamic matrix and newly opened space due to degradation [24]. It has been previously shown that degradable materials favor a spread cell morphology [26] and MSC osteogenic differentiation [27]. The traction forces and integrin-mediated cell interactions in degradable materials are regulated through mechanosensing pathways. Yes-associated protein (YAP) and transcriptional coactivator with PDZ-binding motif (TAZ) represents one of such pathways and enables nuclear mediators of mechanical signals exerted by the ECM and cytoskeletal tension [28]. The activation of this pathway is linked to the integrins on the cell surface and the associated anchoring points in the ECM (e.g. binding motifs) [29]. This, in turn, is modulated by the dimensionality and the degradability of the ECM [30]. YAP/TAZ expression and nuclear translocation as a consequence of cytoskeletal tension, and its effect on cell differentiation [31], cell communication [32] or cell polarization through spatial heterogeneity in YAP activation [33] are well known. However, the influence of multicompartment materials in YAP-mediated mechanotransduction requires further research.

Here, we investigate how 3D patterns in alginate hydrogel degradation affect hMSC behavior and how multicompartment materials differ from single-phase materials. In particular, we induce hMSC spreading, collective alignment, enhanced YAP nuclear translocation and osteogenic differentiation solely with anisotropic hydrogel degradation and under standard cell growth media. Such effect is enhanced under osteogenic media. Additionally, we explore the mechanisms driving osteogenesis in 3D patterned materials, particularly YAP nuclear translocation, and show that both matrix adhesion and degradation characteristics are a necessary requirement. These results advance our understanding of cell response in anisotropic microenvironments. Mimicking tissue anisotropy in 3D patterned hydrogels could have broad applications in biofabrication and tissue engineering.

## 2 Materials and Methods

### 2.1 Alginate modification

Norbornene and tetrazine were added to the alginate backbone via carbodiimide chemistry (see Fig. 1A and further details in Suppl. Fig. 1). The alginate used was low molecular weight, high guluronic acid sodium alginate (MW 75 kDa Pronova UP VLVG; NovaMatrix). The coupling of norbornene (N, TCI Chemicals, #N0907) and tetrazine (T, Conju-probe, #CP-6021) to the alginate molecule was performed as previously described [23]. Alginate modification with norbornene was performed with a theoretical degree of substitution (DS_theo_) of 500. Alginate modification with tetrazine was performed with a DS_theo_ of 170 (Suppl. Table S1). NMR spectroscopy was used to determine the actual DS (DS_actual_) (Suppl. Fig. S2). The quantification of the DS was performed by averaging the characteristic peak area: for tetrazine 10.2, 8.2 and 7.4 ppm and for norbornene 6.2-5.8 ppm, then dividing this value by the area of the characteristic alginate peak (G-1) between 5.2-4.8 ppm. NMR measurements were performed, using a 1.5 % w/v alginate solution in deuterium oxide (64 scans; Agilent 400 MHz Premium COMPACT equipped with Agilent OneNMR Probe) and analyzed using MestreNova Software (version 12.03).

**Figure 1:**
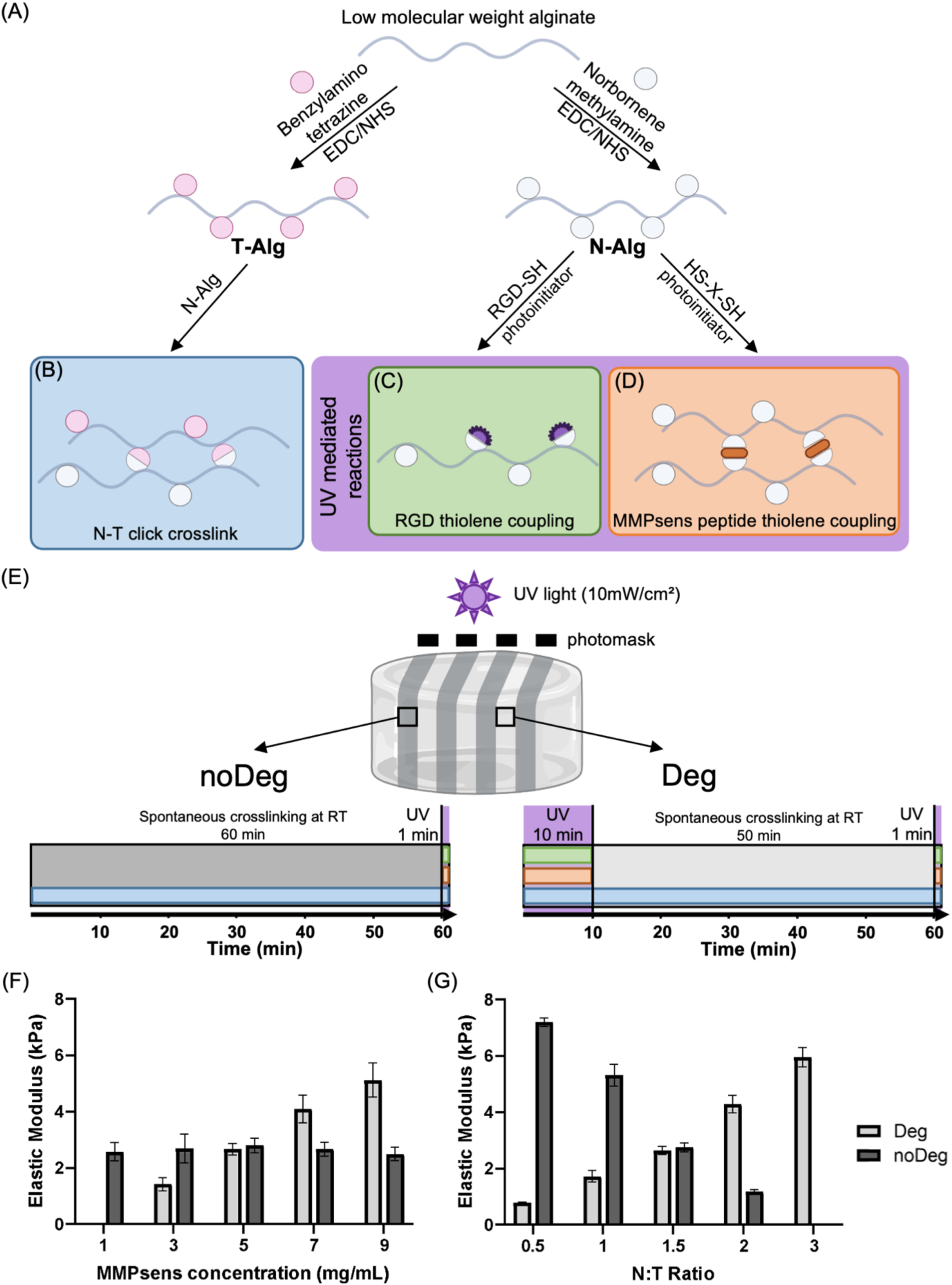
Description of the material chemistry. Scheme of the carbodiimide coupling of norbornene or tetrazine groups to the alginate backbone (A). Scheme of non-degradable bonds through Diels-Alder spontaneous click crosslinking of norbornene and tetrazine (B, blue). UV light dependent thiol-ene reactions of norbornene groups with RGD (C, green) and MMPsens peptide crosslinker (X: GCRD-VPMS ↓ MRGG-DRCG) (D, orange). Time scheme depicting the different reactions occurring during the dual crosslinking (E). Material elastic modulus can be tuned through the MMPsens concentration, for a constant N:T ratio of 1.5 (F) and also through varying the N:T ratio, for a constant MMPsens concentration of 5 mg/mL (G).

### 2.2 Isolation and culture of primary human bone-marrow derived mesenchymal stromal cells (hMSCs)

Bone-marrow derived hMSCs (female, 50 years, no secondary diagnosis, no co-medication) were received from the Core-Facility “Cell Harvesting” of the Berlin Institute of Health (BIH). The cells were isolated from metaphyseal bone marrow biopsies from patients undergoing hip replacement at the Charité - Universitätsmedizin Berlin, as previously described [34][35]. Written informed consent was given, and ethics approval was obtained from the local ethics committee/institutional review board (approval number EA2/089/20) of the Charité – Universitätsmedizin Berlin.

For differentiation analyses, hMSCs in passage P5 were encapsulated in hydrogels and incubated in growth or osteogenic media. For further information on cell isolation and cell culture procedures, please refer to Supplementary Information 1.

### 2.3 Fabrication of anisotropic matrices: 3D patterns in alginate hydrogel degradation

Hydrogels were fabricated based on previously established protocols [23]. The single-phase materials (no photo mask) were used as controls for mechanical testing of bulk properties by rheology with a frequency sweep test (Section 2.4.1), and for cell stainings. In addition, 3D patterned hydrogels were generated as described below. For all material samples, cell culture was performed either in growth or osteogenic.

#### 2.3.1 Non-degradable matrix: Diels-Alder click-crosslinking

The precursors for the hydrogel were dissolved in phosphate-buffered saline (Merck #D8537) and distributed into 2 tubes. The first tube contained norbornene-modified alginate (N-alg); MMP-sensitive (MMPsens) peptide (GCRD-VPMS ↓ MRGG-DRCG, 95 % purity; WatsonBio) at a final concentration of 5 mg/ml of hydrogel, thiolated RGD-peptide (CGGGGRGDSP; Peptide2.0) at a theoretical concentration of 5 molecules of RGD per alginate chain (DS 5, 1.2×10^-3^ mM), and the cell suspension at final concentration of 2×10^6^ cells/mL of hydrogel. The second tube contained tetrazine-modified alginate (T-alg) and the photoinitiator (Irgacure 2959; Sigma-Aldrich, #410896) at a final concentration of 3 mg/mL of hydrogel. The total final concentration of alginate was 2 % w/v at a N:T ratio of 1.5. Despite MMPsens and the photoinitiator being present, these were not activated due to the lack of UV exposure. Nevertheless, the MMPsens and photoinitiator need to be present to allow for patterned materials (see Section 2.3.3).

The two solutions were mixed by pipetting and cast onto a bottom glass plate (borosilicate glass, Filafarm, #1000185), with the casting area being restricted on three sides by glass spacers (Marienfield, #1100420), and immediately covered with a glass slide previously treated with SigmaCoat (≥ 99.5 %; Sigma-Aldrich, #SL2) to prevent adhesion. The gel height was constrained to 2 mm by the thickness of the glass spacers. Spontaneous click-crosslinking for 60 min at room temperature (RT) and in the dark allowed Diels-Alder crosslinking and the N:T covalent bonds to form (Fig. 1B).

In order to ensure a homogeneous binding of the RGD-peptide, crosslinked gels were subsequently exposed to 2 min UV light (365 nm) at 10 mW/cm^2^ (Omnicure S2000) in a custom-built exposure chamber. The RGD-peptide binds to the free norbornenes via thiol-ene reaction (Fig. 1C). The cylindrical hydrogels were punched from the cast gel sheet using 5 mm biopsy punches (Integra Miltex) and incubated in cell culture media at 37 °C and 5 % CO_2_. Further information on the reaction timeline can be found in Fig. 1E.

#### 2.3.2 Degradable matrix: UV-mediated thiol-ene crosslinking with MMPsens peptide

The production of degradable materials followed the same procedure as described in Section 2.3.1, with an additional step for the MMPsens peptide crosslinking. After casting the hydrogel solution between the glass plates, the material was immediately exposed to UV light at 10 mW/cm^2^ for 10 min to initiate the coupling of the degradable MMPsens peptide to the norbornene-modified alginate via thiol-ene crosslinking (Fig. 1D). After the UV exposure, the materials were placed for an additional 50 min at RT in the dark to allow for the N:T covalent bonds to be formed. To ensure a homogenous binding of RGD, the hydrogels were finally exposed again to UV for 2 min (Fig. 1C). Further information on the reaction timeline can be found in Fig. 1E. Hydrogels were punched out and incubated in cell culture media at 37 °C and 5 % CO_2_.

The mechanical properties (measurement method described in Section 2.4) can be modulated by varying the concentration of MMPsens peptide (Fig. 1F) and by tuning the N:T ratio (Fig. 1G). For a constant N:T ratio of 1.5, the non-UV exposed noDeg regions have a constant elastic modulus, while the UV exposed Deg regions results in stiffer matrices with increasing MMPsens peptide concentration (Fig. 1F). In contrast, for constant MMPsens peptide concentration of 5 mg/mL, the non-UV exposed noDeg regions have a decreasing elastic modulus with increasing N:T ratio, while the UV exposed Deg regions results in stiffer matrices with increasing N:T ratio, due to the greater abundance of N-alg available for the thiol-ene crosslinking (Fig. 1G)

#### 2.3.3 Anisotropic matrix: Photolithography of patterned dual crosslinked hydrogels

The creation of patterned materials followed the same procedure as described in Section 2.3.2, with the addition of a photomask placed on top of the cover glass during the UV mediated thiol-ene coupling of the MMPsens peptide. The photomask had a pattern of straight lines with 500 μm thickness (UV light blocking sections, non-degradable matrix equivalent to 2.3.1) placed 300 μm, 500 μm or 800 μm apart (UV light permitting sections, degradable matrix equivalent to 2.3.2). After the 10 min UV exposure through the photomask, followed by 50 min incubation in the dark at RT, in which all the N-T bonds were formed and no additional grafting of MMP occurred, crosslinked gels were exposed to 2 min UV light at 10 mW/cm^2^ without a photomask for homogeneous binding of the RGD-peptide (Fig. 1E).

Two additional control materials are included in this manuscript: materials with a non-degradable peptide sequence and materials with no RGD adhesion peptide. The control materials with a non-degradable peptide sequence were achieved by replacing the MMPsens with the homologous scramble (MMPscr) peptide (GCRD-VpMS ↓ mRGG-DRCG, 95 % purity; WatsonBio), where the amino acids in lower case were switched for the D-isoform, unrecognizable by MMP enzymes. For the control materials with no adhesion peptides, the thiolated RGD solution was replaced with PBS. All of the controls follow the procedure established under Section 2.3.3.

### 2.4 Hydrogel mechanical characterization over time

Mechanical characterization was performed on day 1 and day 14. All mechanical characterization was performed with cell-laden materials to quantify the effect of cell-driven enzymatic degradation of the hydrogels. The material degradation and changes in mechanical properties were evaluated via two different methods: (i) rheology with a frequency sweep test to measure the bulk shear modulus, *G* (and then calculate the elastic modulus, *E*), (ii) microindentation to measure the surface elastic modulus of single-phase and patterned materials, and display the iso-or anisotropic values with a surface map.

#### 2.4.1 Rheology

Storage and loss modulus of single-phase hydrogels were determined with a rheometer (Anton Paar MCR301) via frequency sweeps with a parallel plate geometry of 8 mm (PP08, Anton Paar). The frequency sweep was performed from 0.01 to 10 Hz and at 0.1 % shear strain at RT (n = 6). Once contact with the gel surface was established, a pre-compression of 10 % of the height of the hydrogel was applied prior to the measurement. No additional hydration was needed as the experiment lasted less than 10 min. To obtain the elastic modulus, first the shear modulus (G) was derived from the storage (G’) and loss (G”) modulus using Rubber’s elasticity theory 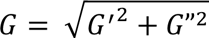 [36]. The elastic modulus (E) was calculated using the values of *G* and the approximation of Poisson’s ratio (𝜗) equal to 0.5, *E* = 2*G* (1 + 𝜗) [37]. The mesh size (𝜉) was approximated based on storage modulus G’ in low frequencies 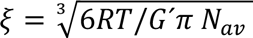, as previously described [38].

#### 2.4.2 Microindentation

To verify the emerging patterns in mechanical properties of the 3D hydrogels due to anisotropic degradation, the surface elastic modulus of the materials on day 1 and day 14 was mapped into a heatmap. Depth-sensing microindentation measurements were done using a Triboindenter TI-950 (Hysitron-Bruker, MN, USA) with a cono-spherical tip of 50 µm radius. The tip was calibrated by an air indent, followed by a quick approach to detect the surface. Similar procedures as previously described were used [23], but were adapted for softer materials. Then, the measurements were automatized as follows: ∼350 µm retraction, an indent approach of ∼150 µm and strain rate of ∼30 µm/s. The measurements were done in automated mode to map a 2 x 2 mm^2^ area with 10 x 10 indents and 200 µm spacing. To ensure that the sample remained hydrated during the measurement, the samples were fixed to a sponge and partially submerged in PBS.

To obtain the elastic modulus from the load-displacement indentation curves, the Hertzian contact model was used. The analysis of the load-displacement curves was automated by a custom-made Python3 script. We used an adapted version of the script described in our previous work [23], related to the selection of the origin point. The origin point is the initial contact point where the tip gets in contact with the material surface and from which the Hertzian model will be determined. This procedure was previously automatized by mathematically determining the minimum of the first part of the indentation curve. However, in very soft materials like the ones presented in this study, the origin point could not be identified using the previous method, as the initial contact point was not as clear as with stiffer gels. The adapted version of the script allows the user to select the origin point manually.

### 2.5 Characterizing volume and wet weight changes over time

To track the degradation of single-phase materials, volume changes, wet weight changes and swelling ratios (S_t_) were tracked over time, at 0 (after 2 h equilibration), 1, 3, 5, 7, 14, 21 and 28 days. At each time point, single-phase hydrogels were weighed under sterile conditions and measured using a caliper. The volume was determined assuming a cylindrical shape, calculated from one height measurement and the mean of two diameter measurements. Swelling (*S*) was calculated with the equation *S_t_=W_t_/W_t=0_*, where W_t_ refers to the hydrogel wet weight at a given time point *t*, and W_t=0_ refers to the hydrogel wet weight at the initial time point.

### 2.6 Primary hMSC viability and cell number

To verify cell viability Live/Dead stainings were performed after 1 and 14 days. The hydrogels were taken out from the incubation media and washed with PBS (Merck #D8537). Then the cells were stained with a solution of 4 µM Calcein AM (Biomol Bioquest #ABD-22002, 500x solution in DMSO) and 4 µM Ethidium homodimer-1 (Sigma #E1903, dilution 1:1000 from original solution) dissolved in PBS to identify live and dead cells, respectively. The staining solution volume was 400 µl per hydrogel in a 24 well plate, stained for 12 min in a cell culture incubator at 37 °C, 5 % CO_2_ in the dark. A final washing step was performed with 400 µl of PBS per hydrogel at RT for 5 min and protected from light.

Imaging was performed on a confocal microscope (Leica SP5, Germany). Quantification of cell number and viability at each time point was performed using ImageJ software (ImageJ 1.53s) [39]. One position from 3 independent samples was acquired at the gel center at 10x water-immersed objective (HCX APO L U-V-I 10.0x 0.30), with tile merging of 3×3 and a 250 µm z-stack.

### 2.7 Primary hMSC morphology

Cell morphology was assessed after 1 and 14 days, staining the nuclei and actin cytoskeleton using SYTOX and Phalloidin, respectively. All steps were performed under orbital shaking in a 24 well plate and using a volume of 500 µL per gel. The culture media was removed and the gels were washed twice with PBS (Merck #D8537). Then, the encapsulated cells were fixed in 4 % paraformaldehyde solution (Sigma-Aldrich, #158127) for 45 min at RT, followed by permeabilization with 0.3 % Triton X-100 (Sigma-Aldrich, #11488696) for 15 min, washed twice with 3 % bovine serum albumin (BSA, Sigma-Aldrich, #A7906) in PBS for 5 min. The staining solution was dissolved in PBS containing 0.1 % TritonX-100 and 3 % BSA and with the following concentrations: 0.5 µM SYTOX (Thermo Fisher, #S11381) and 1 µg/mL of Phalloidin 405 Conjugate (CruzFluor™, #SC363790). The staining was performed protected from light, during 4 h at RT. Final washing was performed twice with 3 % BSA in PBS for 5 min at RT.

For imaging, three different positions per gel were acquired at the gel center from two independent samples using a confocal microscope (Leica SP5, Germany). For a general quantification of cell morphology, 10x water-immersed objective (HCX APO L U-V-I 10.0×0.30), with a tile merging of 2×2 was used and a 250 µm z-stack.

### 2.8 Primary hMSC differentiation

Cell differentiation was tested using Osterix (OSX) as an early osteogenic differentiation (see section 2.9), Osteocalcin (OCN) as medium osteogenic differentiation (see section 2.9), and OsteoImage as late osteogenic differentiation marker. The stainings were performed after 14 days, always in combination with cell morphology staining, SYTOX and Phalloidin, as described in Section 2.7.

For OsteoImage (Lonza, #PA1503) the staining procedure followed the provideŕs instructions. Briefly, the fixed cells (see Section 2.7) were washed with the wash buffer twice for 5 min. Then, the previously diluted 1:100 Osteoimage staining reagent in dilution buffer was added to the gels, 500 µL per gel. The incubation was carried out at 4 °C, overnight and protected from light. The next day, the samples were washed three times with the wash buffer and then imaged.

For imaging, three different positions per gel were acquired at the gel center from two independent samples using a confocal microscope (Leica SP5, Germany). For a higher magnification quantification of cell morphology, 25x water-immersed objective (HCX IRAPO L 25.0×0.95), with a tile merging of 4×4 was used and a 250 µm z-stack.

### 2.9 Immunostainings

YAP, Osterix (OSX) and Osteocalcin (OCN) expression was evaluated by immunofluorescence. After performing the morphological staining, SYTOX and Phalloidin, as described in Section 2.7, the primary antibody was incubated for 24 h at 4 ⁰C under constant rocking. The antibodies used were: YAP IgG2aκ mouse monoclonal (Santa Cruz, #sc-101199) at 10 µg/mL in PBS, OSX polyclonal rabbit (abcam, ab22552) 10 µg/mL in PBS-1%BSA and OCN monoclonal mouse IgG1 (R&D Systems, #MAB1419) at 25 µg/mL in PBS. After the primary antibody staining, the samples were washed three times with 400 µL of 3 % BSA in PBS for 30 min. The secondary antibody (ThermoFisher, #A32728) was stained for 2 h at RT under constant rocking. Finally, the samples were washed 3 times with 3 % BSA in PBS and imaged.

For imaging, three independent positions per gel were acquired at the gel center from two independent samples using a confocal microscope (Leica SP5, Germany). For a higher magnification quantification of cell morphology, 25x water-immersed objective (HCX IRAPO L 25.0×0.95), with a tile merging of 4×4 was used and a z-stack of 250 µm.

### 2.10 Quantitative analysis of stained sections

The cell morphology in 3D patterned materials was evaluated using a previously described macro [23] written in ImageJ (ImageJ 1.53s, [39]). The cell projected area, cell circularity and cell number were obtained from z-stack projections. The macro offers the possibility to freely divide an image into a grid, which leads to a heat map in which the results are later depicted. The cell border threshold was manually set.

To evaluate collective cell alignment, the Image J plugin Directionality was used [40]. The graphs show the ‘Direction (°)’ or angle vs. the ‘Amount’, which shows the fraction of cells with a preferred direction, normalized to the total cell number.

YAP, Osterix (OSX) and Osteocalcin (OCN) expression was quantified as a percentage of the marker expressing cells (YAP, OSX or OCN or positive cells) divided by the total cell number (SYTOX).

The images were evaluated using ImageJ: first, a background subtraction of a control image of the staining with only the secondary antibody was performed, followed by setting manually the threshold and finally evaluating with the automatic particle counting function. The mineralized area obtained with OsteoImage (OI) staining, was established as a percentage by dividing the OI positive cell area by the total cell area (phalloidin staining).

The nuclear to cytoplasmic ratio of YAP was evaluated using a previously described method [41]. Briefly, individual cells were segmented for nucleus (SYTOX) and cytoplasm (phalloidin) to define the respective subcellular regions. Next, YAP average intensity within each subcellular region was quantified and the ratio calculated.

### 2.11 Flow cytometry analysis

After 14 days of encapsulation, the cells were retrieved from the hydrogels using alginate lyase (Sigma, #A1603, reconstituted in Ca/Mg-free PBS in 2mg/mL). The cells were incubated for 30 min at 37C, then centrifuged at 400 rcf during 5 min and washed twice with PBS. The flow cytometry staining and analysis was done as described before [42]. The viable cells were determined using fixable live/dead blue staining kit for UV excitation (Thermo Fisher L34962). The blocking solution (FcX, BioLegend, 422301) was used to prevent unspecific binding. Staining with antibodies (Supplementary table 2) was performed on ice and using flow cytometry buffer (phosphate-buffered saline (PBS) including bovine serum albumin (0.5% w/v) and sodium acid (0.1% w/v)). Fixation and permeabilization kit (BioLegend, 426803) was used for intracellular staining. The stained cells were analyzed on an ID7000 spectral flow cytometry (Sony Biotechnology, San Jose, CA, USA) with individual autofluorescence unmixing. Gating and analysis of the population were performed with FlowJo software (Tree Star, USA), further analysis were performed in R Statistical Software (v4.4.1; R Core Team 2024) using the Spectre packages [43].

### 2.12 Statistical analysis

Statistical analysis was performed with GraphPad Prism 8 (version 8.4.3 for Windows, GraphPad Software Inc., Boston, Massachusetts USA). To test for normality, the Shapiro-Wilk test was applied. For normally distributed data, bar plots showing mean and standard deviation (SD) were used and the statistical comparison was performed using ANOVA (* alpha < 0.05, ** alpha<0.01 or *** alpha<0.001), corrected with Bonferroni test for multiple comparisons. For not normally distributed data, box plots with median, 1^st^ and 3^rd^ quartile were used and data was compared using a Kruskal-Wallis test (* alpha < 0.05, ** alpha<0.01 or *** alpha<0.001), corrected with Dunńs test for multiple comparisons.

## 3 Results

### 3.1 Anisotropic degradation of cell-laden patterned hydrogels leads to emerging patterns in mechanics

Cells were encapsulated in photopatterned alginate hydrogels (Fig. 2A) with initial homogeneous mechanical properties (Fig. 2J). The comparable mechanical characteristics of the materials were achieved by tuning the MMPsens peptide concentration (Fig. 1F) and the N:T ratios (Fig 1G). The mechanical properties of cell-laden hydrogels evolved during time due to the cellular enzymatic action (Fig. 2A). Those mechanical properties were characterized by rheology (Fig. 2B, C, D, G) and microindentation (Fig. 2H-J).

**Figure 2:**
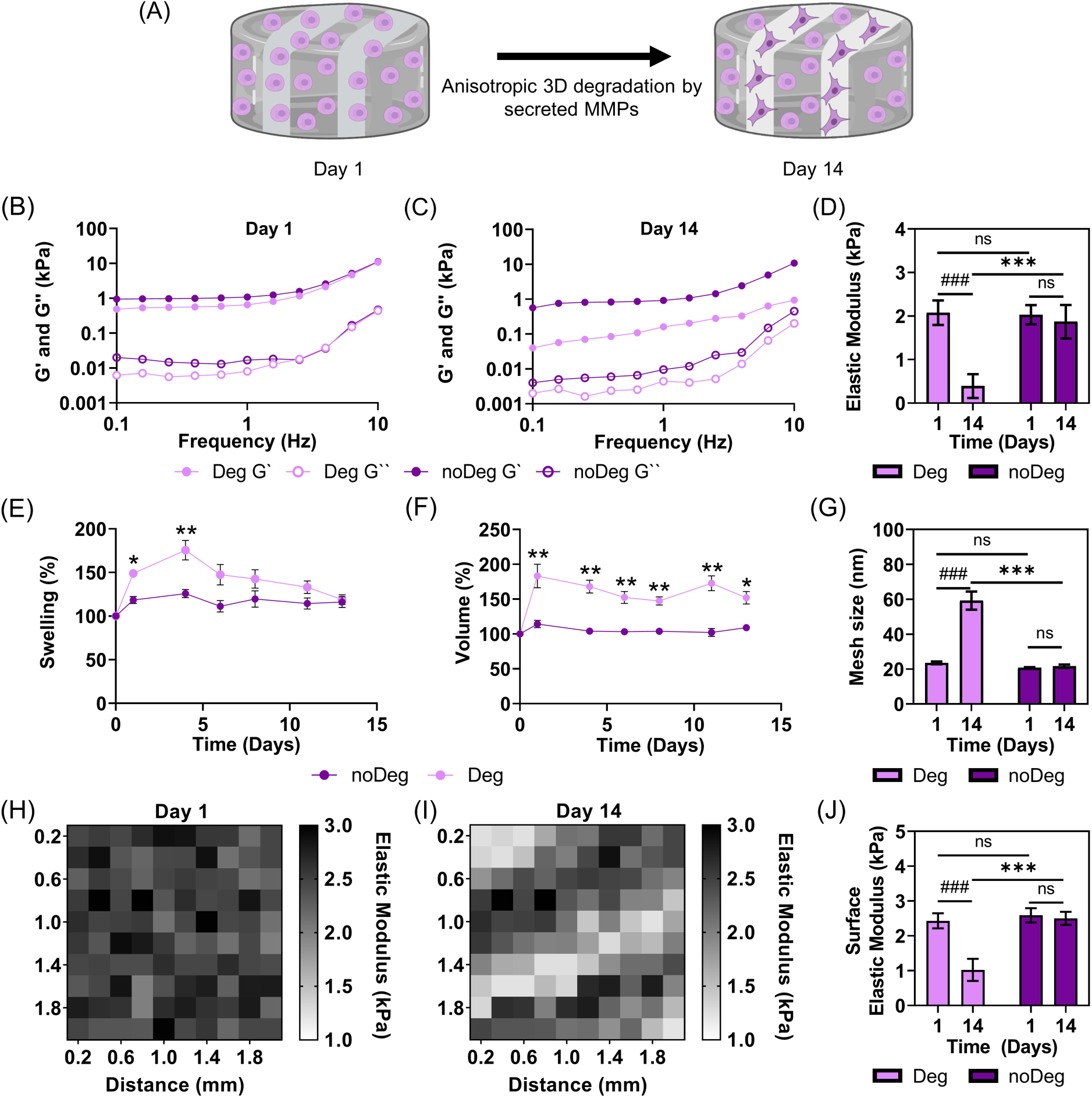
Changes in mechanical properties and swelling over time of cell-laden anisotropic hydrogels. Anisotropic dual crosslinking results in 3D patterns of hydrogel degradation due to cell-derived enzymatic action (A). Representative plots of G’ (storage modulus, ●) and G’’ (loss modulus, ○) obtained by rheology of noDeg single-phase (dark purple) and Deg single-phase (light purple) materials, on day 1 (B) and day 14 (C) from n = 6 gels. Elastic modulus (D) and corresponding mesh size (G), approximated from rheology data. Swelling (E) and volume (F) changes over time (relative to 2h after equilibration), from n = 3 gels. Representative microindentation map of the surface elastic modulus on day 1 (H) and day 14 (I), 10×10 grid, indentation spacing of 200 µm. Surface elastic modulus in Deg and noDeg phases obtained by microindentation, averaging all indents in a specific region of patterned materials on day 1 and day 14 (J). Statistical significance with ANOVA and Bonferroni correction.

The degradable and non-degradable materials have comparable values of loss (G’) and storage (G’’) modulus on day 1 (Fig. 2B). However, on day 14 there is a pronounced difference in G’ and G’’ for Deg materials (Fig. 2C). A significant decrease over time in mechanical properties of Deg materials is observed, whereas the noDeg materials preserve their elastic modulus (Fig. 2D). The elastic modulus on day 1 of Deg (2.07 ± 0.28 kPa) is comparable to noDeg (2.03 ± 0.22 kPa). Significant differences can be seen between the elastic modulus of Deg (0.39 ± 0.27 kPa) and noDeg (1.87 ± 0.38 kPa) on day 14. The elastic modulus changes can be related to the water content as there is a significant difference in the swelling (Fig. 2E) and volume changes over time (Fig. 2F), which is also reflected in an increasing mesh size (Fig. 2G). The swelling and volume of Deg gels starts to decrease after 5 days due to degradation, while a less dynamic behaviour is seen in the noDeg materials. The mesh size of Deg materials increases significantly between day 1 (24 ± 1 nm) and day 14 (59 ± 5 nm), whereas in noDeg materials the mesh size remains unchanged from day 1 (21 ± 0.3 nm) to day 14 (22 ± 1 nm).

Compared to the degradation behavior of single-phase materials, the corresponding phases in patterned materials show similar characteristics, as shown by the average surface elastic modulus of microindentation measurements (Fig. 2J). As seen in the representative surface map of elastic modulus, the pattern in degradation emerges over time. There is no distinct pattern of the surface elastic modulus on day 1 (Fig. 2H), with values fluctuating around 2.5 kPa, consistent with the comparable bulk elastic modulus of both single-phase materials (Fig. 2D). However, on day 14 (Fig. 2I), there are softer sections (1-1.5 kPa) corresponding to the Deg zones and stiffer sections (2.5 kPa) associated with noDeg zones.

### 3.2 Anisotropic hMSC morphology and collective cell alignment in 3D patterned materials compared to single-phase counterparts

hMSCs were encapsulated in 3D anisotropic materials with patterns in degradation. The viability and cell number were evaluated using Calcein/ EtHD1 staining. Cell viability (fraction of Calcein (+) cells) remained high on day 1 (Suppl. Fig. S3A-B) and day 14 (Suppl. Fig. S3D-E) and there were no visible patterns in cell viability. Cell proliferation was also evaluated by individual quantification of the average cell number (total cells, Calcein and EtHD1) (Suppl. Fig. S3C and S3F).

Next, we investigated the effect of patterned hydrogels on cell morphology and collective alignment. Patterned materials with dimensions of 500 µm:500 µm for Deg:noDeg (Fig. 3A) were compared on day 14 to single-phase materials (Fig. 3E, 3G). On day 1 (Suppl. Fig. S4A-C), cell morphology was similar among the two phases of the patterned material, not showing any significant differences, neither in projected cell area (Fig. 3H) nor in circularity (Fig 3I). Patterns in cell morphology were visible on day 14 (Fig. 3A) and were quantified through heatmaps (Fig 3B-C). The results showed increased projected cell area (Fig. 3B) and decreased circularity (Fig. 3C) in Deg regions, while cells remained round and with smaller projected cell area in noDeg zones. Similar trends were observed in equivalent single-phase materials (Fig. 3D-G). However, the effect in the morphological changes was significantly enhanced in the Deg sections of patterned materials in contrast to the single-phase Deg (Fig. 3H-I).

**Figure 3:**
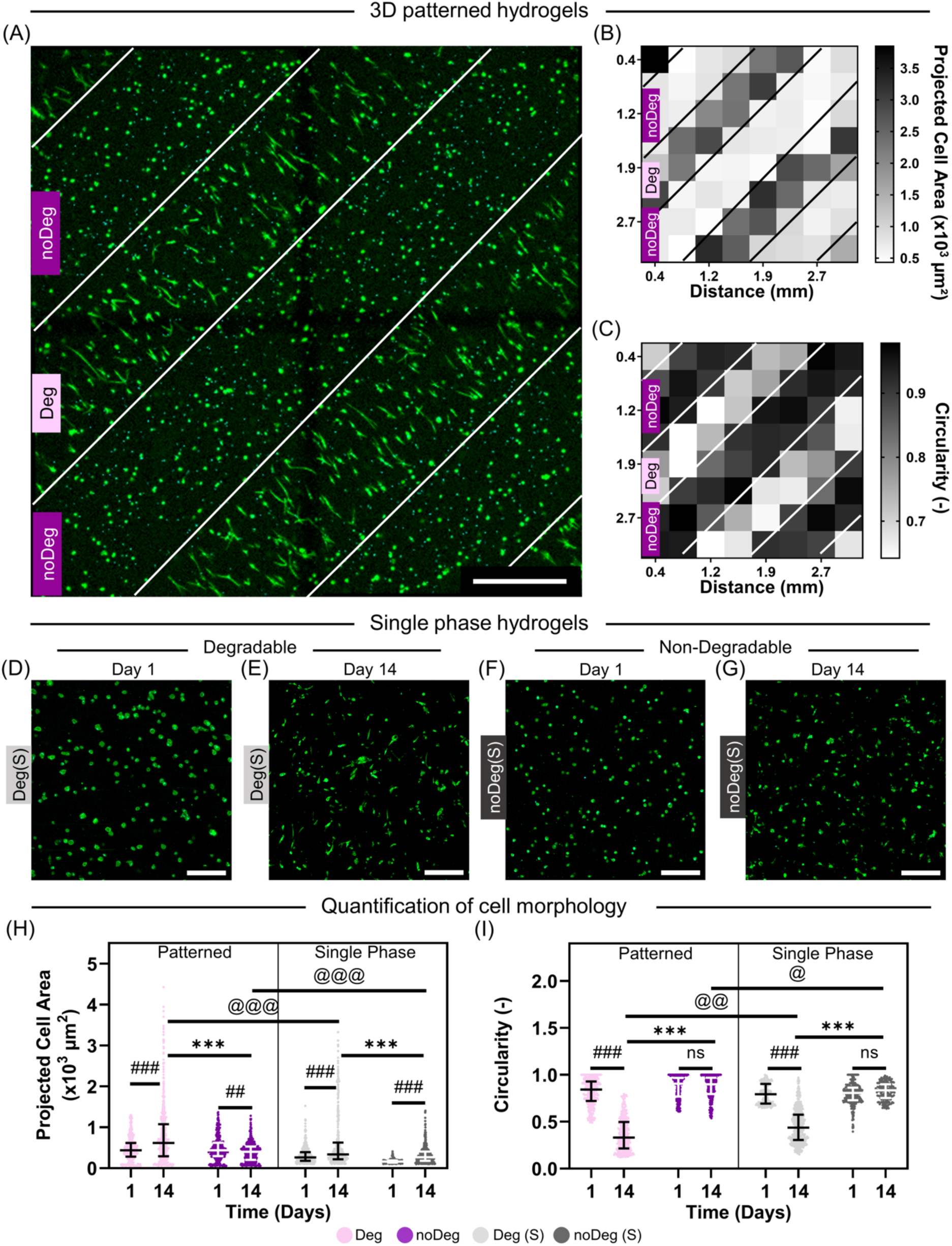
Anisotropic hMSC morphology in patterned materials on day 14. Representative confocal images of actin (Phalloidin/green) and nucleus (SYTOX/cyan) staining of patterned materials with hMSCs in growth media (A). Heatmap quantification with an 8×8 grid of projected cell area (B) and cell circularity (C). Single-phase materials representative confocal images of actin (Phalloidin/green) and nucleus (SYTOX/cyan) staining on day 1 (Deg (D) and noDeg (F)) and day 14 (Deg (E) and noDeg (G)). Dot graph of raw data (> 400 single cells per phase and condition) of projected cell area (H) and circularity (I) on day 14 (see Suppl. Fig. S4A-C for corresponding images and heat maps on day 1). Statistical significance with Kruskal-Wallis and Dunn’s correction). Scale bar: 500 µm (A) and 250 µm (D-G).

The patterned materials allowed enhanced changes in cell morphology but also differences in collective cell alignment (Fig. 4). The cell orientation was evaluated for patterns of different Deg:noDeg dimensions, 300 μm:300 μm (Fig. 4A-C), 500 μm:500 μm (Fig. 4D-F) and 800 μm:200 μm (Fig. 4G-I). The 300 μm:300 μm photomask (Fig 4A) resulted in cells collectively aligned perpendicular to the stripes in the Deg phases (Fig 4B), in contrast to the cells in noDeg sections (Fig 4C), where cells remained round and had no predominant orientation. This effect was further enhanced with the 500 μm:500 μm photomask (Fig. 4D-F). However, for the 800 μm:200 μm photomask (Fig 4G), with a broader degradable section, the collective cell alignment was lost (Fig. 4H) and was equivalent to single-phase materials (Suppl. Fig. 5A-D), where we found a spread cell morphology but lacked a collective cell alignment. Similarly, no collective alignment was found in patterned materials when subjected to constrained swelling in the XY direction (Suppl.Fig.5E-L).

**Figure 4:**
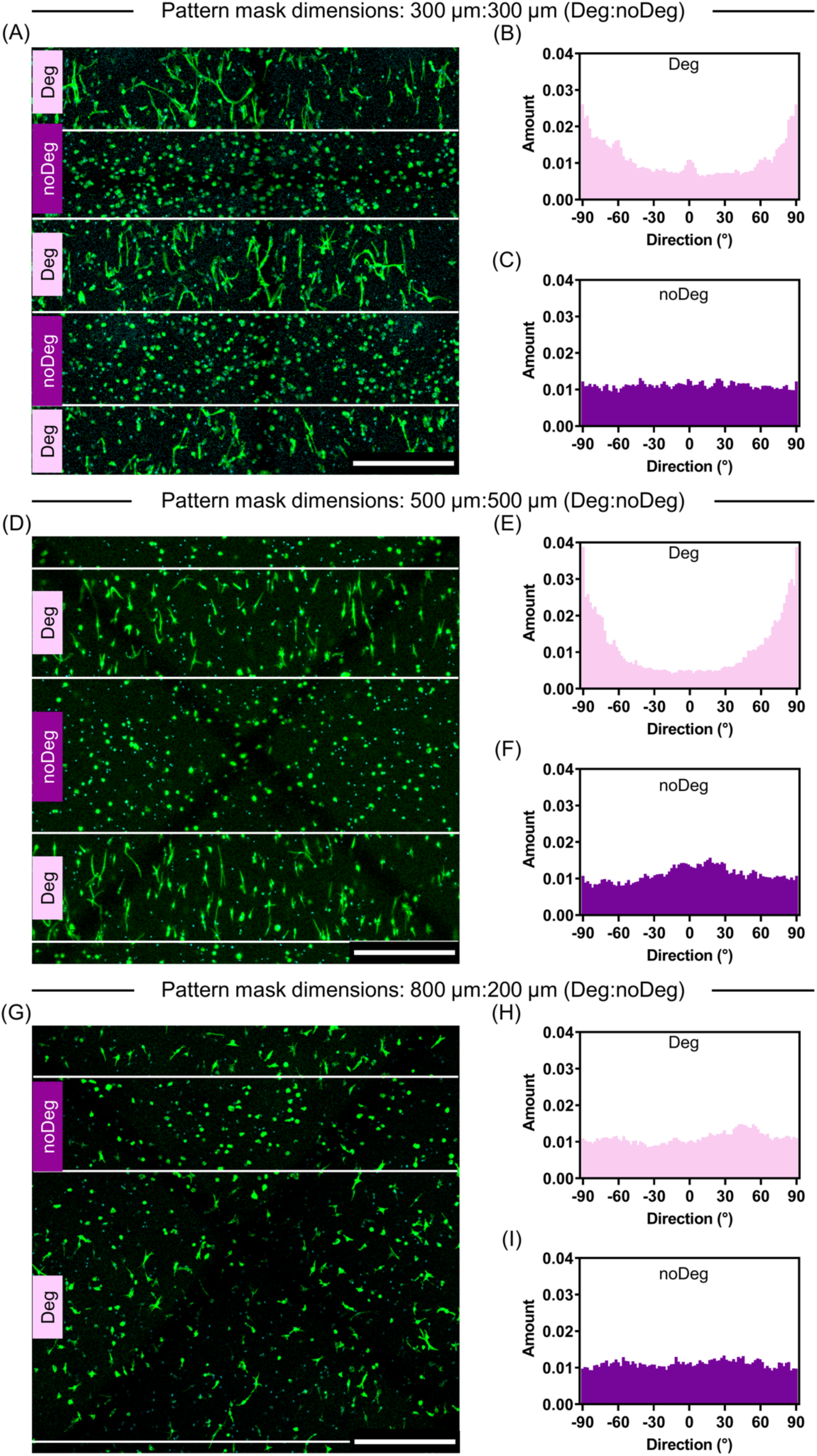
Collective cell alignment of encapsulated hMSCs in patterned materials with different photomask dimensions on day 14. Patterned materials fabricated with photomasks of different Deg:noDeg dimensions: 300 μm:300 μm (A-C), 500 μm:500 μm (D-F) and 800 μm:200 μm (G-I). Representative confocal images of actin (Phalloidin/green) and nucleus (SYTOX, cyan) staining of patterned materials with hMSCs (A, D, G) and histograms indicating the distribution of the collective cell alignment angle in Deg and noDeg phases of patterned materials (B/C, E/F, H/I), with n > 200 cells per phase. Scale bar: 500 µm.

**Figure 5:**
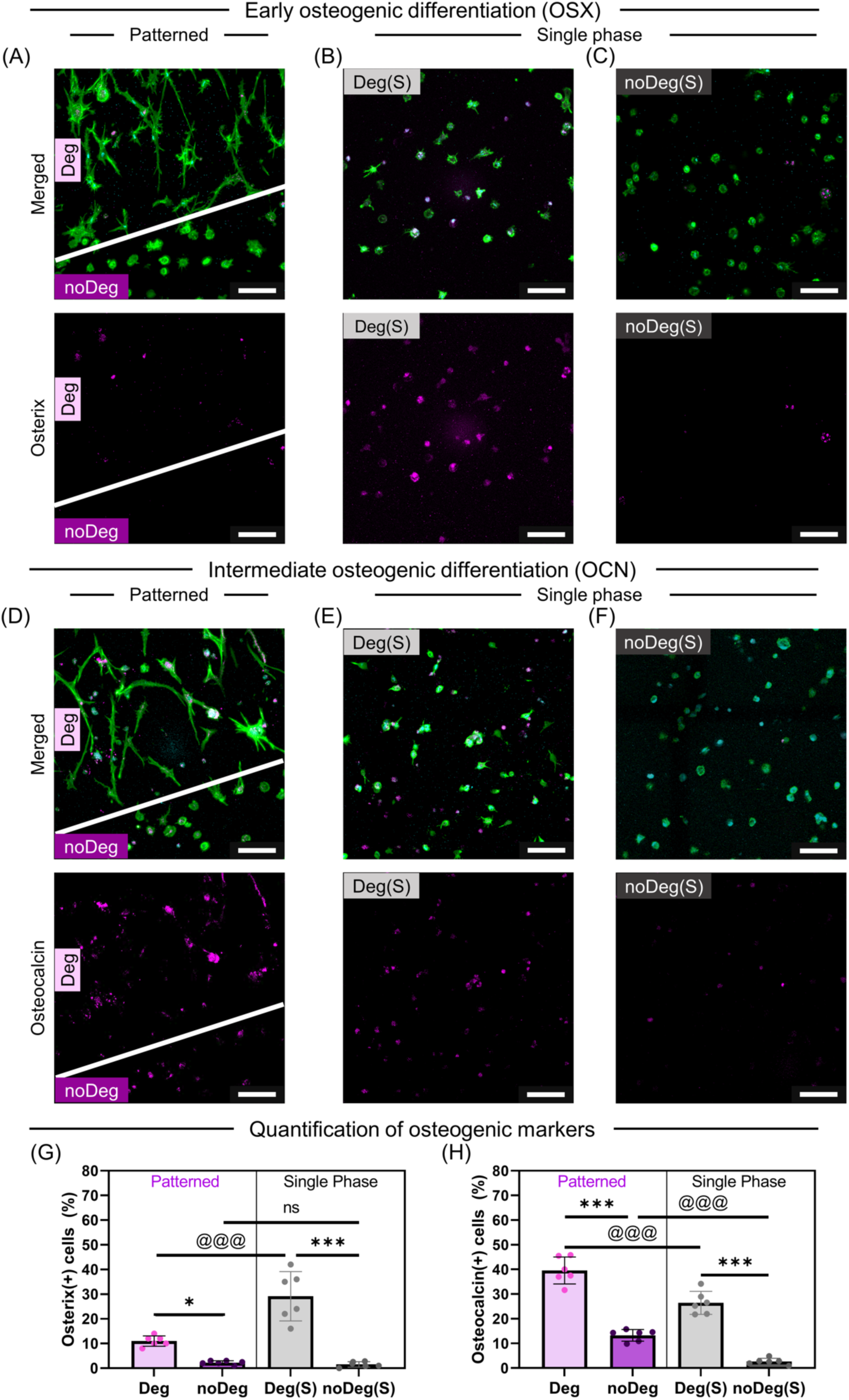
Osteogenic differentiation of encapsulated hMSCs in patterned materials on day 14. Actin (Phalloidin/green) and nucleus (SYTOX/cyan) staining with Osterix (OSX, magenta, A-C) or Osteocalcin (OCN, magenta, D-F). Comparison between patterned materials (A, D), single-phase Deg(S) (B, E) and single-phase noDeg(S) (C, F), quantified in bar plots (G, H), which show the mean and standard deviation of n = 6 fields of view containing multiple cells (n ≥ 150 cells per phase and condition). Statistical significance with ANOVA and Bonferroni correction. Scale bar: 100 µm.

### 3.3 Enhanced and spatially guided hMSC osteogenic differentiation in 3D patterned materials compared to single-phase counterparts

In the previous sections we showed that 3D patterns in alginate hydrogel degradation induced an anisotropic response in hMSC morphology and emergent properties such as collective cell alignment. This motivated further investigation of the anisotropic hMSC osteogenic differentiation, evaluated by early (Osterix, OSX) and intermediate (Osteocalcin, OCN) osteogenic markers.

The anisotropic biophysical cues provided by the patterned materials already induced significantly enhanced and spatially segregated osteogenic differentiation of hMSC over 14 days in growth media (Fig. 5). The early osteogenic marker OSX (Fig 5A-C) was expressed in the Deg single-phase materials (Fig. 5B) and also in the Deg phase of patterned materials (Fig. 5A), though to a lower extent (Fig. 5G). Furthermore, the intermediate osteogenic marker OCN, indicative of further differentiated hMSCs, was significantly higher (Fig. 5H) in the Deg sections of patterned materials (Fig 5D) compared to single-phase Deg materials (Fig. 5E). The noDeg phases in both patterned and single-phase materials (Fig. 5C, F) did not allow osteogenic differentiation of hMSCs.

With flow cytometry, individual cells can be explored on a higher parametric level. Cells were retrieved from three different types of hydrogels: single-phase degradable (Deg(S)), single-phase non-degradable (noDeg(S)), and patterned 3D hydrogels (patterned), after culturing for 14 days. We investigated the changes in cell phenotype by analyzing various cell surface markers, as well as intracellular markers, allowing to trace the differentiation from stem-like characteristics towards osteogenic cells (Fig. 6). Dimension reduction (UMAP) of all cells from the three groups showed distinct clustering of the groups (Fig 6A-G). The mean fluorescence intensity (geometric MFI) allowed to determine the expression level of measured markers. All retrieved cells from the 3D materials were negative for the markers CD45, CD34, CD166, CD271 and CD146. The surface markers CD90, CD105, CD73, considered as minimal criteria for hMSC, showed altered positive expression levels with CD90 (Fig 6H), especially, being higher expressed in the degradable compared to the noDeg material (Fig 6H, K).

**Figure 6:**
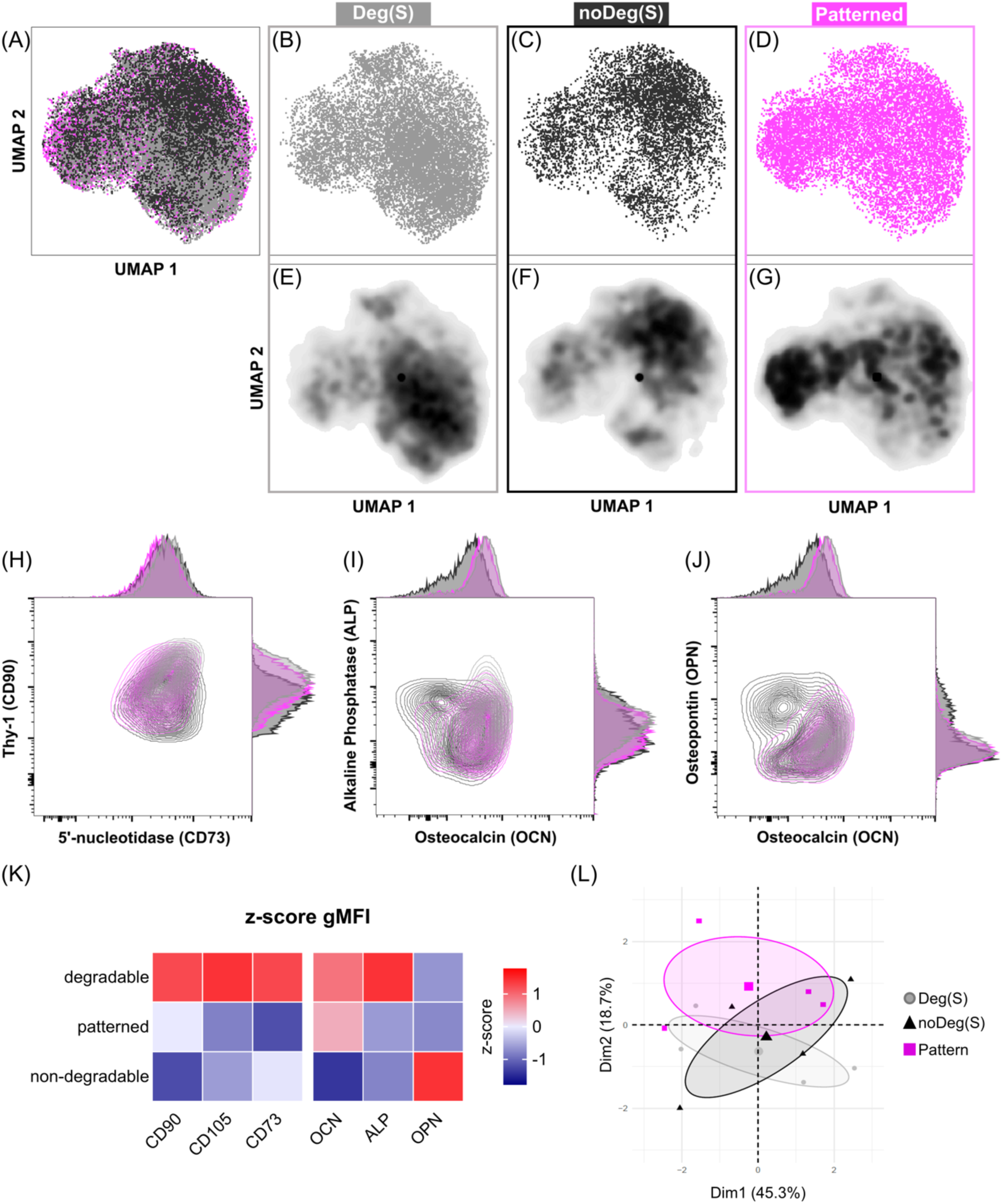
Flow cytometry evaluation of hMSCs retrieved from 3D patterned and single-phase materials after culturing for 14 days. UMAPs cluster plots merged (A), Deg(S) (B), noDeg(S) (C) and patterned (D), and the corresponding density plots of Deg(S) (E), noDeg(S) (F) and patterned (G). Multiplots, contour plots plus histograms of the expression of: CD73 vs. CD90 (H), OCN vs. ALP (I) and OCN vs. OPN (J). z-score quantification of hMSCs minimal criteria markers (CD105, CD73, CD90) and differentiation markers (OCN, ALP, OPN) (K). Principal component analysis for dimension 1 (Dim 1, 45,3 %) and dimension 2 (Dim 2, 18,7 %), the big symbols show the center of the ellipse and the smaller symbols the different batches (L). n = 4 batches containing 1×10^5^ cells, for each material.

The intracellular markers Osteocalcin (OCN), Alkaline Phosphatase (ALP) and Osteopontin (OPN) allowed to trace the osteogenic differentiation of encapsulated hMSCs, which show an increased z-score of OCN and ALP in Deg materials (Fig. 6I, K). Interestingly, a small subset of cells in the noDeg material expressed higher levels of OPN (Fig 6J, K).

To evaluate the difference between the patterned and single-phase materials, a principal component analysis was performed (Fig 6L). The x-axis represents Principal Component 1 (PC1), which explains 45.3% of the variance, while the y-axis represents Principal Component 2 (PC2), capturing 18.7% of the variance. Despite the overlap between the groups, the trend shows 3 distinct groups enclosed in a 90% CI ellipse with an overall 64% explained variances. This indicates that the patterned material induced a distinct cellular composition that differs from the respective single-phase materials.

### 3.4 Enhanced and spatially guided mechanosensing via YAP/TAZ signaling in 3D patterned materials compared to single-phase counterparts

After establishing the significant differences in patterned hMSC projected cell area, circularity, collective cell alignment and osteogenic differentiation, we investigated the underlying mechanism. Based on the observation of morphological changes and associated cytoskeletal tension in degradable regions, we investigated YAP positive expression (YAP(+)) as a binary measurement, independent of intensity, as well as its nuclear to cytoplasmic localization within single cells in patterned materials (Fig. 7A-C) and compared it to the single-phase counterparts (Fig 7D-G).

**Figure 7:**
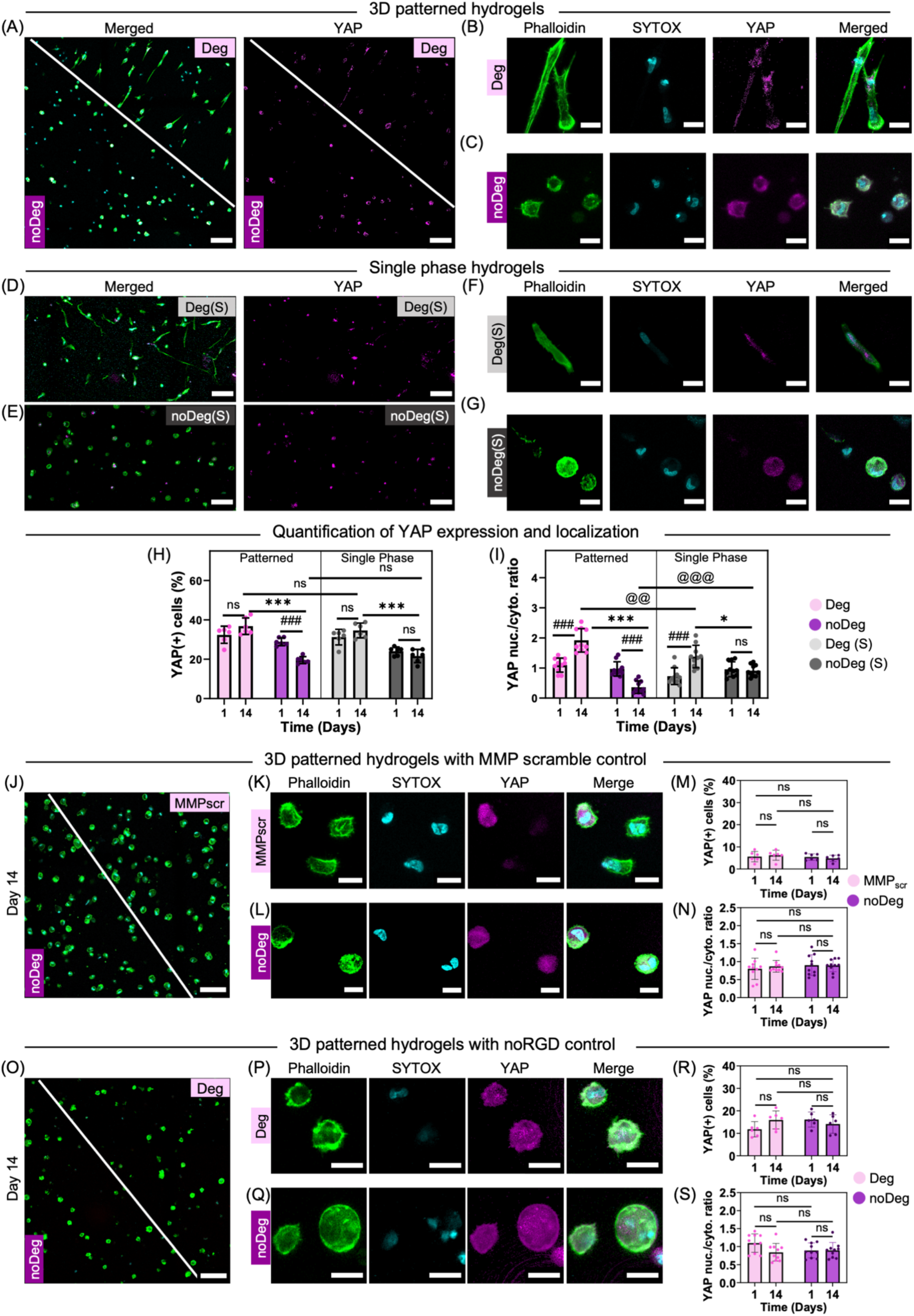
YAP expression and nuclear to cytoplasmic localization of encapsulated hMSCs in patterned materials vs. single-phase materials, and controls with MMP scramble peptide and without RGD on day 14. hMSC staining with actin (Phalloidin/green), nucleus (SYTOX/cyan) and YAP expression (magenta). YAP expression in patterned materials: overview (A), representative single cell images in the Deg (B) and noDeg (C) phase. YAP expression in single-phase materials: overview of Deg (D) and noDeg (E) and representative single cells in Deg (F) and noDeg (G) materials. Comparison between patterned and single-phase materials of YAP expression (H) and nuclear to cytoplasmic localization (I). YAP expression in patterned materials with MMPscr control: overview (J) and representative single cell images in Deg (K) and noDeg (L) regions. Quantification of YAP expression (M) and YAP localization (N) in MMPscr materials. YAP expression in patterned materials with noRGD control: overview (O) and representative single cell images in Deg (P) and noDeg (Q) regions. Quantification of YAP expression (R) and YAP localization (S) in noRGD materials. Bar plots show the mean and standard deviation of n = 6 fields of view containing n > 120 cells (H, M, R) and of n = 10 representative single cells from Deg vs. noDeg regions (I, N, S). Statistical significance with ANOVA and Bonferroni correction. Scale bar: 100 µm (A, D, E, J, O) and 20 µm (B, C, F, G, K, L, P, Q).

The fraction of YAP(+) cells was significantly higher in Deg materials patterned (Fig 7A) and single-phase (Fig 7D) compared to noDeg (Fig. 7E) on day 14 (Fig 7H). Independent of the material or the patterning, the YAP expression was equally distributed over the nucleus and the cytoplasm on day 1 (Fig 7I). On day 14, there was a nuclear translocation (nuc./cyto. >1) in Deg materials, patterned (Fig 7B) and single-phase Deg (Fig 7F), whereas for the noDeg materials, in patterned (Fig 7C) and in single-phase (Fig 7G), it remained close to 1. The effect of nuclear YAP translocation was significantly higher in Deg phases of patterned materials compared to Deg phases in single-phase materials (Fig. 7I).

To test the influence of degradation in the YAP translocation, the MMPsens peptide crosslinker was replaced with the homologous non-degradable scramble peptide (MMPscr). The MMPscr allowed the comparison of the degradable and non-degradable material with the same chemical crosslinking and mechanical properties (ex. crosslinker length). The MMPscr patterned materials showed a very similar behavior on day 1 (data not shown) compared to day 14 (Fig. 7J-L). The lack of degradation in the MMPscr regions led to a rounded cell morphology with a similar YAP expression (Fig. 7J-K) as the one seen in noDeg phases (Fig. 7G). Both phases with MMPscr and noDeg patterned materials had a similar YAP (+) cell percentage (Fig. 7M) and YAP localization (Fig. 7N), with no significant differences neither between phases nor over time.

To test the relevance of integrin activation and cell-matrix interaction with alginate, we used materials lacking RGD adhesion peptides (Fig. 7O-S). The cells in noRGD patterned materials showed a similar behavior in the two phases on day 1 (data not shown) as well as on day14 (Fig 7O-Q). The fraction of YAP(+) cells was lower (Fig. 7R) compared to materials with RGD (Fig. 7H). Moreover, there were no significant differences in YAP(+) cells (Fig 7R) or localization (Fig. 7S), neither between phases nor over time.

### 3.5 Additional biochemical stimuli can enhance the spatially guided hMSC osteogenic differentiation

hMSC differentiation was further tested under osteogenic media to evaluate the effect of biochemical stimuli in patterned materials. Cell morphology patterns are enhanced with osteogenic media (Fig. 8A-E) when compared to growth media (Fig. 3A), showing larger projected cell area (Fig. 8B) and elongated cells with lower circularity (Fig. 8C) in Deg regions.

**Figure 8:**
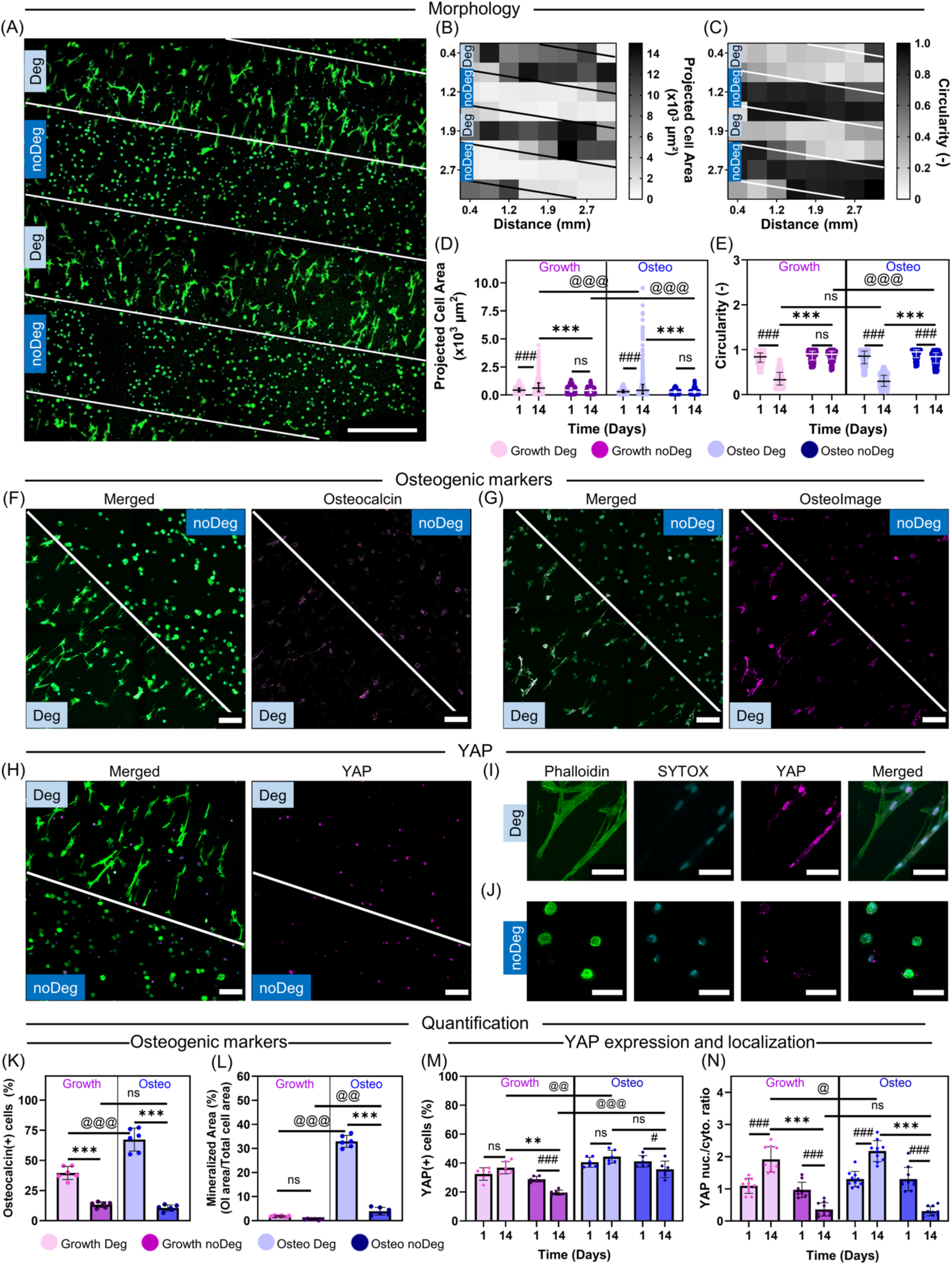
Enhanced cell response in patterned materials by osteogenic media on day 14. Representative actin (Phalloidin/green) and nucleus (SYTOX/cyan) staining of patterned materials with hMSCs in osteogenic media (A). Heatmap quantification with an 8×8 grid of projected cell area (B) and cell circularity (C). Dot graph of growth vs osteogenic media of cells in patterned materials (> 400 single cells per phase and condition) of projected cell area (D) and circularity (E) on day 14 (see Suppl. Fig. S4D-F for corresponding images and heat maps on day 1). Representative Osteocalcin (magenta, F) or Osterix (magenta, G) staining and expression quantification of OCN (K) and mineralization (L). Representative YAP staining overview (H) and single cells from Deg (I) and noDeg (J) sections. Quantification fraction of YAP(+) cells (M) and YAP localization (N). Dot graphs show statistical significance with Kruskal-Wallis and Dunn’s correction and bar plots evaluated with statistical significance with ANOVA and Bonferroni correction. Scale bar: 500 µm (A), 250 µm (F, G, H) and 20 µm (I, J).

The intermediate osteogenic marker OCN had higher expression in osteogenic media (Fig. 8F) compared to growth media and shows a significantly higher expression in Deg phases compared to noDeg phases (Fig. 8F, K). Despite the biochemical stimuli of osteogenic media, the restrictive environment of noDeg sections resulted in only a low number of OCN positive cells (Fig 8K). The late osteogenic marker (mineralization, OsteoImage) (Fig. 8G, L) also had higher expression in osteogenic media compared to growth media, with significantly higher mineralized area in Deg sections compared to noDeg sections (Fig. 8L).

The fraction of YAP(+) cells in osteogenic media (Fig. 8H) showed also increased expression on day 14 in the Deg regions, but with overall increased values compared to growth media (Fig. 5M). The YAP nuclear to cytoplasmic ratio shows a similar significant pattern over time in osteogenic media compared to growth media (Fig. 8N), with nuclear translocation in the Deg phase (Fig. 8I) and cytoplasmic expression in the noDeg phase (Fig. 8J). In the Deg phase the use of osteogenic media significantly enhanced the nuclear translocation compared to growth media (Fig. 8M, N).

## 4 Discussion

Degradation patterned materials enhance encapsulated hMSC spread morphology, allow collective cell alignment, stimulate mechanotransduction and promote osteogenic differentiation, when compared to single phase materials. By investigating patterned hydrogels as opposed to equivalent single-phase matrices, we aimed to highlight the importance of using multicompartment materials in biomaterial design. In addition, the patterned materials with anisotropic hydrogel degradation allowed elucidating the role of degradation, without additional biophysical (such as stiffness) or biochemical (such as osteogenic media) cues. Our findings offer insights into how these anisotropic materials influence cell response, potentially leading to better strategies mimicking tissue anisotropy, with applications in biofabrication and tissue engineering.

The anisotropic alginate-based hydrogels used in this research consist of two soft (<3kPa) 3D microenvironments for cells, with comparable initial elastic modulus at day 1, and emerging patterns through degradation over time. The comparable initial elastic modulus of both phases allows a direct comparison on degradability effects, excluding other biophysical properties (i.e. matrix stiffness), which have already been studied [44][24]. Moreover, the combination of both matrices in a patterned multicomponent material results in a structural advantage over time, as the single phase Deg materials crumble and start disintegrating due to degradation, while patterned materials keep their shape after 14 days (Suppl. Fig. S6). It is important to highlight the tuneability of the matrix platform used, which allows different combinations of mechanical and degradation properties (Fig. 1F-G). Interestingly, the click reaction described in Fig. 1G shows a maximum elastic modulus at N:T ratio 0.5. This is consistent with the description of an unexpected dimer formation, following a 1:2 (N:T) stoichiometry [45].

Cell spreading is dependent on the matrix degradation and this is enhanced in Deg sections of patterned materials compared to single phase counterparts. Changes in morphology are a result of the remodeling of the immediate environment [30] and consistently described in MMP-sensitive peptide crosslinked materials [46]. Our research showed that the interphases of the patterned materials not only lead to spatially segregated cell morphology, but also influence the collective cell alignment perpendicular to the stripe main axis. Cell alignment in patterned materials has been previously described on 2D topographical directed patterns [47][48][49] and in 2D mesh scaffolds [50], exhibiting a parallel alignment to the groves/ridges, or perpendicular to them in wider curvatures (>1000 µm) [51]. However, less is known about the mechanism for collective cell alignment within 3D matrices. Previous studies report the perpendicular cell alignment to be a result of force and neighboring tensile stress [52]. Our research showed that cell alignment is dependent on the stripe dimensions (Fig 4, broader stripes show no alignment), patterning (Suppl. Fig S5, single phase hydrogels show no collective alignment) and, probably, on the materiaĺs differential swelling properties (Suppl. Fig S5, constrained swelling show no alignment). The direct transfer of mechanisms of collective cell alignment into 3D matrices is still at early stage, as 3D environments offer multiple factors that can influence cell, such as mechanical properties, adhesive ligand concentration, crosslink type and density. Yet, understanding and guiding collective cell alignment will be important for developing functional biomaterials that mimic biological anisotropic tissues, which rely on fiber/cell directionality for optimal performance [53].

The 3D patterns in hydrogel degradation allowed spatially guided hMSC differentiation, promoting enhanced osteogenic differentiation in the Deg region compared to the noDeg, or single-phase degradable materials in growth media (Fig. 5). Strikingly, an early osteogenesis stage as shown by OSX expression could be promoted solely by biophysical factors after 14 day culture in growth media (hMSCs in Deg single-phase materials, Fig. 5B). This effect was enhanced in patterned materials, which at the same time point showed OCN expression, an intermediate osteogenic marker (Fig 5D). Our findings agree with previous studies on stiffness-degradation 3D patterned materials showing that soft and Deg matrices foster osteogenesis [24]. Our study highlights the importance of matrix degradability for osteogenesis, and the potential of using patterned materials for enhancing and guiding osteogenic differentiation.

Moreover, biochemical factors (osteogenic media) in combination with patterned materials, enabled late osteogenesis and mineralization in Deg phases. Notably, mineralization within the Deg phases was only observed under osteogenic media, consistent with studies showing that biomineralization is a complex process [54]. Despite the biochemical cues from osteogenic media were available for cells in both Deg and noDeg phases of the patterned material, the guidance of the patterns is evident with broader cell spreading, higher osteogenic differentiation and increased YAP nuclear translocation in the Deg phases compared to the noDeg regions. These findings highlight the interplay between biochemical and biophysical cues modulating cell behavior and the synergistic effect by anisotropic patterned hydrogels.

Further flow cytometry analysis of hMSCs encapsulated in hydrogels enabled us to identify changes in cell phenotype between the materials. The density plots showed clearly distinct clustering behavior of the Deg and noDeg single-phase materials (Fig 6B, C), however the patterned material did not cluster along the two single-phase materials (Fig 6D, G), hinting towards distinct processes in anisotropic materials and not only replicating the two individual isotropic materials. All cells were consistent with the minimal criteria markers: negative for CD45, CD34, and positive for CD90, CD73, CD105 [55]. There were no hMSCs subpopulations identified as the cells were negative for CD166, CD271 and CD146 [56]. Deg materials led to the upregulation of CD90 (Fig. 6H), which controls vital cellular processes, including cell–cell and cell–matrix interactions, cell adhesion and cell response to dynamic matrices [57][58]. Notably, CD73 shows a downregulation in the patterned materials when compared to Deg and noDeg. This effect could be due to the additional mechanical strain from the anisotropic material interfaces, as previous studies have shown that CD73 is downregulated in mechanically loaded cells [59].

The intracellular markers Osteocalcin (OCN), Alkaline Phosphatase (ALP) and Osteopontin (OPN) allowed to trace the osteogenic differentiation of encapsulated hMSCs. In accordance with the immunofluorescence staining for Osteocalcin (Fig 5), Deg and patterned materials induced osteogenic differentiation in the encapsulated cells, as shown by the increased OCN expression (Fig 6 I, J) compared to noDeg. It is important to highlight that the patterned materials were analyzed as a bulk (Deg+noDeg regions), therefore the enhanced effects in patterned Deg phase were not captured using this method. This is a limitation of bulk analyses using flow cytometry, as opposed to image-based methods and immunofluorescence. Further bulk analyses of cell phenotype, such as RNA seq, were no longer considered, as they would have the same limitations with patterned samples.

Interestingly a small subset of cells in the noDeg material expressed higher levels of OPN (Fig 6 J). OPN is a multifunctional protein, which has a role in inflammatory processes [60]. Non-degradable alginate-based matrices have been previously reported to upregulate differentially expressed genes involved in inflammatory response in breast cancer cells [61].

YAP nuclear translocation and mechanotransduction is enhanced in Deg sections of patterned materials. In line with previous studies, Deg materials showed a higher YAP nuclear translocation compared to noDeg, triggered solely by the degradation of the material [31][30][62]. Our research takes this effect further by evaluating the enhanced mechanotransduction effect in patterned materials compared to single-phase counterparts (Fig 7A-I). The enhanced hMSCs spreading and collective cell alignment resulting from the anisotropic material interphases, seem to enhance mechanosensation by YAP nuclear translocation. As underlying mechanism, we identify two key factors: matrix degradation (as no YAP nuclear translocation was observed in MMPscr materials, Fig. 7J-N) and the need of cell-matrix adhesion via integrin activation (as no YAP nuclear translocation was observed in noRGD materials, Fig. 7O-S). This ultimately leads to enhanced mechanotransduction and osteogenic differentiation in anisotropic hydrogels.

The research presented in this paper represents a contribution to the basic understanding of how cells interact with multicompartment materials. In the future, further research will be required to understand the *in vivo* implications of anisotropic materials.

## 5 Conclusions

In summary, we show that anisotropic hydrogel degradation enhances hMSC cell spreading in 3D, collective cell alignment, mechanotransduction and osteogenic differentiation, compared to the single-phase equivalents. Patterned materiaĺs interphase proximity, matrix-bound cell adhesion peptide RGD and degradable crosslinking peptides are required for patterned mechanosensing. These combined results highlight the importance of developing 3D patterned biomaterials to mimic the anisotropy in biological tissues, for both fundamental understanding of guided collective cell behavior and tissue anisotropy, as well as for applications in biofabrication and tissue engineering.

## Acknowledgements

The authors acknowledge the support from all group members of Cipitria, Schmidt-Bleek and Duda’s laboratories. We thank the Cell Core Facility of Charité-Universitätsmedizin for providing the cells used in this study. In addition, we thank the Research Workshop at the Charité-Universitätsmedizin Berlin for developing and manufacturing some experimental devices. We thank Andreas Engels for printing the photomasks used in this research. We thank Mario Thiele from Charité-Universitätsmedizin and Samik Real from Hasso Plattner-Institute for contributing with the macros used in this research. In addition to the support given by Ansgar Petersen and Erik Brauer from JWI.

## 6 CRediT authorship contribution statement

A Cipitria conceived the idea. CA Garrido, A Cipitria developed the methodology. CA Garrido, DS Garske performed the experiments. CH Bucher performed the flow cytometry experiments and data analysis. S Amini supported the micro indentation experiments. CA Garrido quantified and analyzed the data. CA Garrido and A Cipitria drafted the manuscript. DS Garske contributed with the visualization of the data. A Cipitria, K Schmidt-Bleek and GN Duda supervised, reviewed and edited the original data and manuscript. All authors discussed the results and contributed to the final manuscript.

## 9 Data and code availability

All raw and processed data are available in the publicly accessible Edmond repository of the Max Planck Society: https://doi.org/10.17617/3.WLTITB

