## Supplementary information for "Anisotropic Hydrogel Degradation Enhances 3D Collective Mesenchymal Stromal Cell Alignment, Mechanotransduction and Osteogenic Differentiation"

- Supplementary Figure S1: Detailed description of the material chemistry
- Supplementary Figure S2: NMR spectra of modified VLVG alginate
- Supplementary Table S1: Degree of substitution of modified alginate
- Supplementary Information 1: Extended methods on isolation and culture of primary human bone-marrow derived mesenchymal stromal cells (hMSCs)
- Supplementary Information 2: Markers used in flow cytometry
- Supplementary Figure S3: Viability and cell number of encapsulated cells in patterned materials on day 1 and day 14
- Supplementary Figure S4: Cell morphology of encapsulated hMSCs in patterned materials on day 1 in growth and osteogenic media
- Supplementary Figure S5: Cell alignment in single phase materials and constrained swelling patterned materials
- Supplementary Figure S6: Macroscopic pictures of the hydrogels with encapsulated cells 14 days after encapsulation

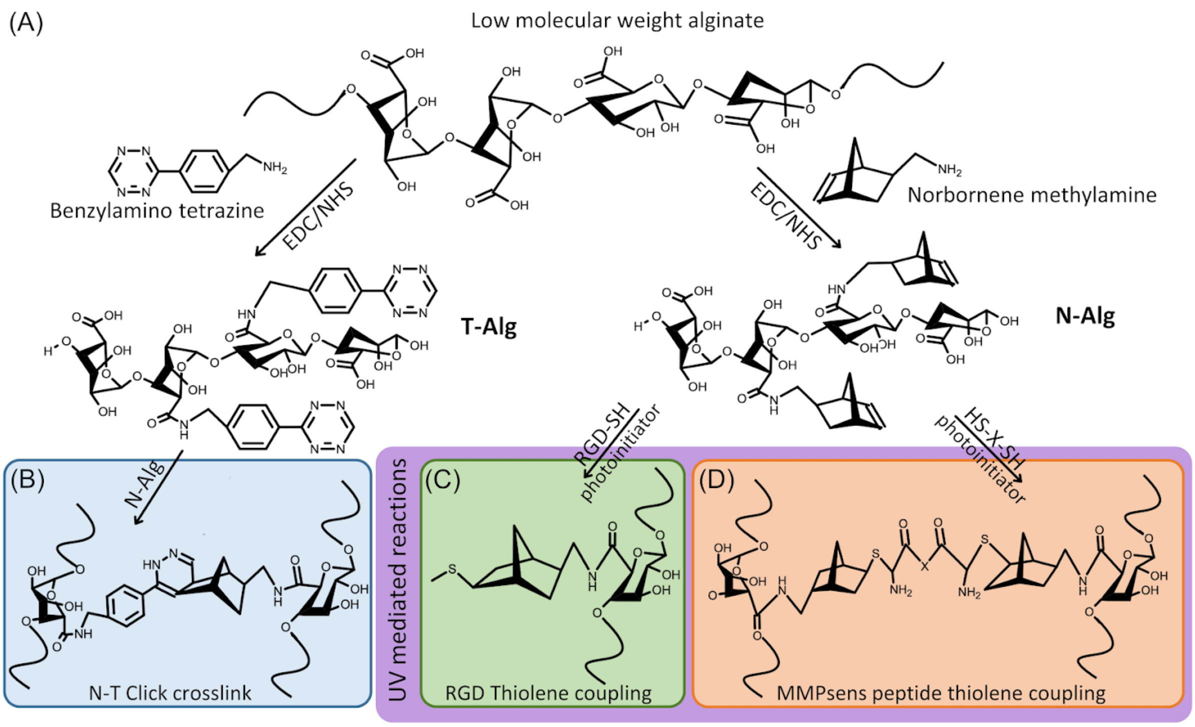


**Supplementary Figure S1:** Detailed description of the material chemistry. Scheme of the carbodiimide coupling of norbornene or tetrazine groups to the alginate backbone through the terminal amine groups to the carboxy groups of the guluronic acid (A). Scheme of non-degradable bonds through Diels-Alder spontaneous click crosslinking of norbornene and tetrazine (B, blue). UV light dependent thiol-ene reactions of norbornene groups with RGD (C, green) and MMPsens peptide crosslinker (X: GCRD-VPMS ↓ MRGG-DRCG) (D, orange).

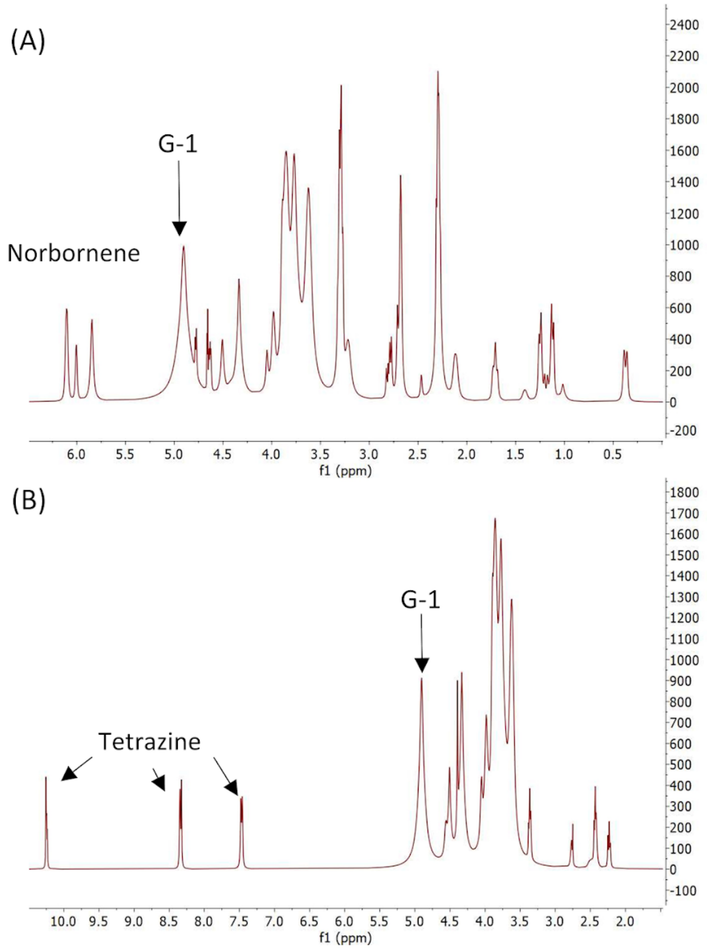


**Supplementary Figure S2: NMR spectra of modified VLVG alginate.** NMR of norbornene modified alginate with the 3 characteristic peaks of norbornene between 6.2-5.8 ppm (A). NMR of tetrazine modified alginate with the 3 characteristic peaks of tetrazine at 10.2, 8.2 and 7.4 ppm (B). In both graphs the H in the first position of the guluronic acid in alginate (G-1), corresponding to the peak between 5.2-4.8 ppm, is used as a reference to determine the $DS_{actual}$.

| | Norbornene | Tetrazine |
|---|---|---|
| $DS_{theo}$ | 500 | 170 |
| $DS_{actual}$ | 43,6 | 36,5 |

**Supplementary Table S1: Degree of substitution of modified alginate.** Norbornene (N-Alg) and tetrazine (T-Alg) modified alginate with a theoretical degree of substitution ($DS_{theo}$) and actual DS ($DS_{actual}$).

**Supplementary Information 1: Extended methods on isolation and culture of primary human bone-marrow derived mesenchymal stromal cells (hMSCs)**

Briefly, the bone marrow mononuclear cell fraction and the stromal cell fraction post Ficoll-density gradient centrifugation (Histopaque 1077; Sigma-Aldrich) were quantified with an automated electrical impedance-based CASY® Cell Counter (Schaerfe System GmbH). The stromal cell-containing interphase was cultured in growth media: Dulbecco's Modified Eagle's Medium (DMEM low glucose, Sigma-Aldrich, #D5546), 10 % v/v fetal bovine serum (Sigma-Aldrich, #F9665), 1 % Glutamax (Gibco, #35050-038) and 1 % penicillin/streptomycin (Gibco, #15140-122). Cells were maintained in a 5 % $CO_2$ environment at 37 °C, medium exchanged every 2 days and passaged when 80 % confluence was achieved. For 3D encapsulation, cells were used at passage 5.

For differentiation analyses, cell-laden hydrogels were incubated in growth or osteogenic or media. Osteogenic media contained: DMEM low glucose (Sigma-Aldrich, D5546) with 10 % v/v fetal bovine serum (Sigma-Aldrich, #F9665), 1 % v/v penicillin-streptomycin (Gibco, #15140-122), 50 µM ascorbic acid (Sigma-Aldrich, A5960), 10 mM ß-glycerolphosphate (Sigma-Aldrich, #G9422) and 100 nM dexamethasone (Sigma-Aldrich, #D4902)

| | Marker | Fluorochrome | Supplier | CatNo | Description |
|---|---|---|---|---|---|
| 1 | CD45 | BUV563 | ThermoFisher | 365-0459-41 | hMSCs minimal criteria, negative markers |
| 2 | CD34 | PE-Cy5 | BD Biosciences | 561819 | |
| 3 | CD105 | BV480 | BD Biosciences | 569877 | hMSCs minimal criteria, positive markers |
| 4 | CD73 | BV605 | BioLegend | 344023 | |
| 5 | CD90 | BV711 | BioLegend | 328139 | |
| 6 | CD166 | BV421 | BD Biosciences | 566280 | hMSCs subpopulation markers |
| 7 | CD271 | APC-Fire750 | BioLegend | 345115 | |
| 8 | CD146 | PE-Cy7 | BioLegend | 361007 | |
| 9 | Osteocalcin | AF647 | NovusBio | IC1419R | Osteogenic differentiation markers |
| 10 | ALP | AF488 | NovusBio | FAB1448G | |
| 11 | Live/Dead | Blue | ThermoFisher | L34962 | Initial cell gating |

**Supplementary Table 2:** Antibodies used for flow cytometry.

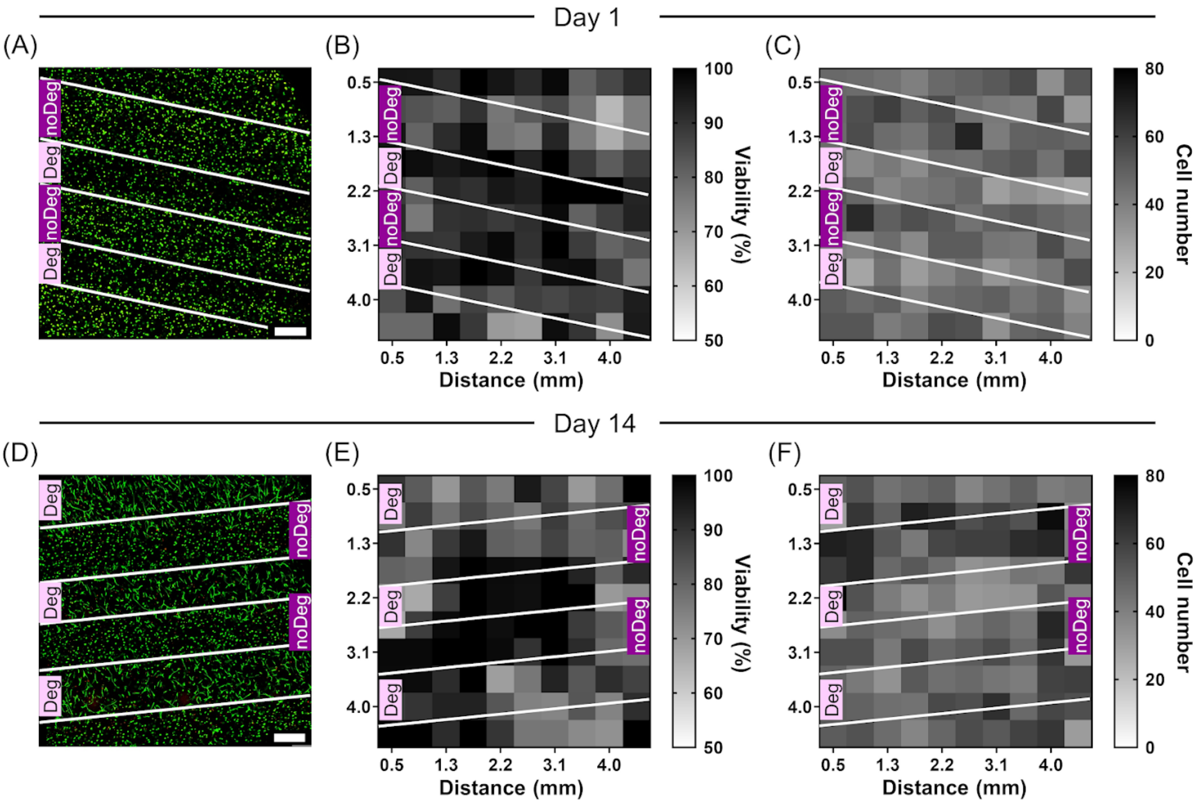


**Supplementary Figure S3: Viability and cell number of encapsulated cells in patterned materials on day 1 and day 14.** Representative confocal images of Calcein (green)/EthD1 (red) staining of hMSCs encapsulated in patterned materials and cultured in growth media on day 1 (A) and day 14 (D). Heatmap quantification of viability (%) with a 10x10 grid on day 1 (B) and day 14 (E). Heatmap quantification of cell number (cells in the field of view) with a 10x10 grid on day 1 (C) and day 14 (F). Scale bar: 500 µm.

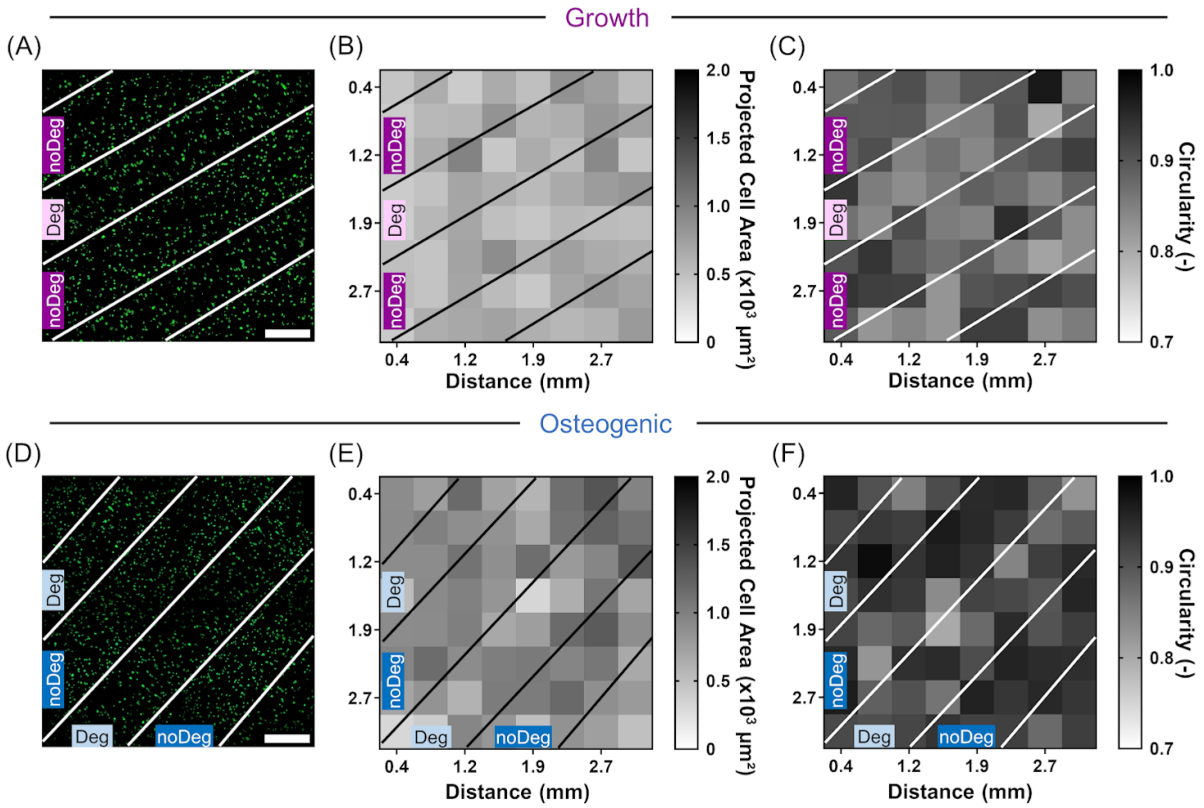


**Supplementary Figure S4: Cell morphology of encapsulated hMSCs in patterned materials on day 1.** Representative confocal images of actin (phalloidin/green) and nucleus (SYTOX/ cyan) staining of patterned materials with hMSCs in growth (A-C) and osteogenic (D-F) media. Heatmap quantification of projected cell area ($\mu m^2$) with an 8x8 grid in growth (B) and osteogenic (E) media. Heatmap quantification of cell circularity with an 8x8 grid in growth (C) and osteogenic (F) media. Scale bar: 500 μm.

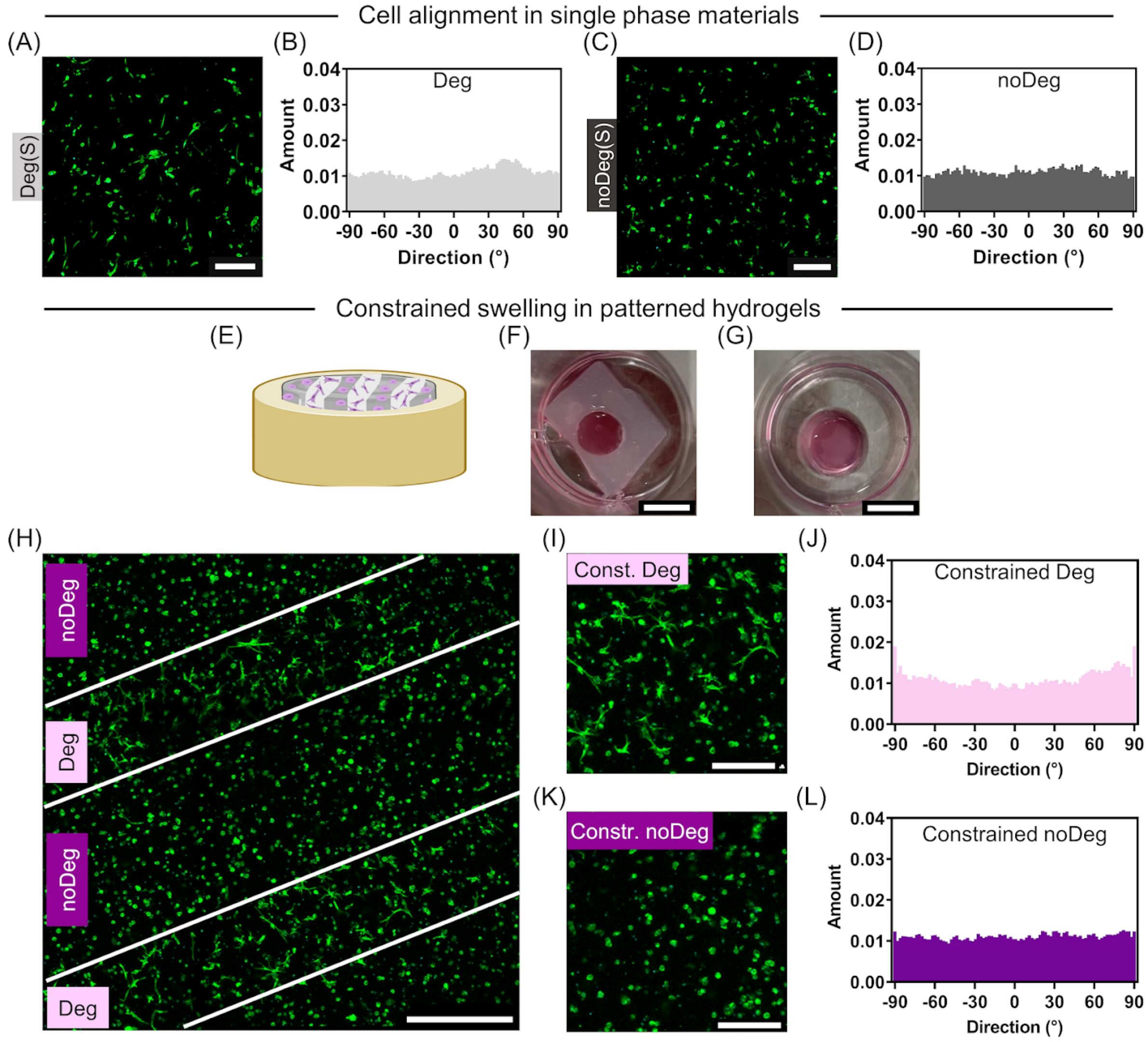


**Supplementary Figure S5: Cell alignment in single phase materials and patterned materials under constrained swelling.** Actin (Phalloidin/green) and nucleus (SYTOX/cyan) staining of encapsulated cells in Deg (A) and noDeg (C) single-phase materials on day 14, and histograms showing the distribution of the collective cell alignment angle in Deg (B) and noDeg (D) materials. Sketch representing the constrained swelling in XY of a patterned material (E). Photos of patterned materials on day 14 under constrained (F) and free (G) swelling. Representative phalloidin/SYTOX staining of a patterned material under constrained swelling on day 14 (H) with a close up of the Deg (I) and noDeg (K) phase. Histograms indicating the distribution of the collective cell alignment angle in Deg and noDeg phases of patterned materials. Scale bar: 500 µm (H), 200 µm (I-K).

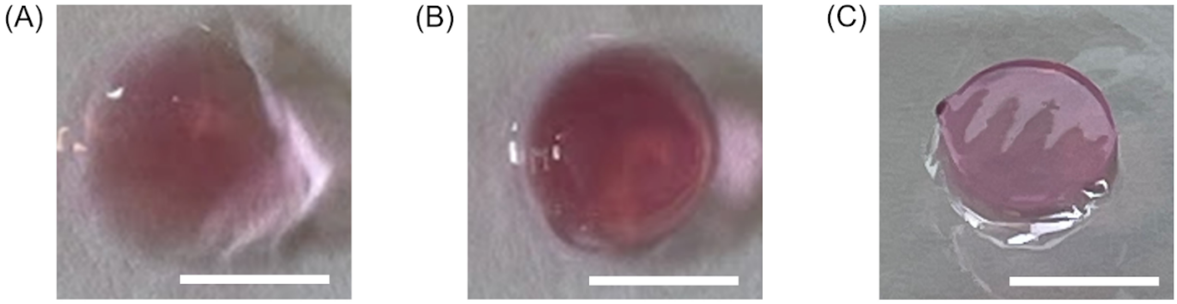


**Supplementary Figure S6: Macroscopic pictures of single-phase and patterned hydrogels with encapsulated hMSCs on day 14.** Single-phase Deg (A), single-phase noDeg (B) and patterned hydrogel (C). Scale bar 5 mm.
